# Perineuronal nets affect memory and learning after synapse withdrawal

**DOI:** 10.1101/2021.04.13.439599

**Authors:** Jiri Ruzicka, Marketa Dalecka, Kristyna Safrankova, Diego Peretti, Giovanna Mallucci, Pavla Jendelova, Jessica CF Kwok, James W Fawcett

## Abstract

Perineuronal nets (PNNs) enwrap mature neurons, playing a role in the control of plasticity and synapse dynamics. PNNs have been shown to have effects on memory formation, retention and extinction in a variety of animal models. It has been proposed that the cavities in PNNs which contain synapses can act as a memory store, which remains stable after events that cause synaptic withdrawal such as anoxia or hibernation. We examine this idea by monitoring positional memory before and after synaptic withdrawal caused by acute hibernation-like state (HLS). Animals lacking hippocampal PNNs due to enzymatic digestion by chondroitinase ABC or knockout of the PNN component aggrecan were compared with wild type controls. HLS-induced synapse withdrawal caused a memory deficit, but not to the level of naïve animals and not worsened by PNN attenuation. After HLS, animals lacking PNNs showed faster relearning. Absence of PNNs affected the restoration of inhibitory and excitatory synapses on PNN-bearing neurons. The results support a role for hippocampal PNNs in learning, but not in long-term memory storage.

## Main

Perineuronal nets (PNNs) are dense extracellular matrix structures surrounding specific neuronal types in the brain and spinal cord. In the brain they are predominantly formed around the fast-spiking parvalbumin positive inhibitory interneurons (PV^+^). They are composed mainly of hyaluronan, chondroitin sulphate proteoglycans (CSPGs), hyaluronan and proteoglycan link proteins (Haplns) and tenascins, the CSPGs being mainly responsible for their inhibitory properties. The individual components can be produced by both glia and neurons(Deepa et al., 2006; Fawcett et al., 2019; Frischknecht et al., 2009; Irvine and Kwok, 2018; Jager et al., 2013; Lensjo et al., 2017a; Ueno et al., 2019). The PNNs form a lattice-like structure on the membrane of PV neurons with a regions of uncovered membrane mostly occupied by synapses (Arnst et al., 2016; Frischknecht et al., 2009). The PNNs are fully formed at the end of critical period and serve as regulators of plasticity and excitability and are also neuroprotective (Beurdeley et al., 2012; Hou et al., 2017; Reichelt et al., 2019). PNNs are modified by behavioural events, and the sulphation pattern of their CSPGs changes with injury and age (Foscarin et al., 2017; Karetko-Sysa et al., 2014; Ueno et al., 2019; Ueno et al., 2018). The number and anatomy of PNNs can change as a result of various behavioural events, during development and aging through regulation on the level of individual PNN components (Gottschling et al., 2019; Saroja et al., 2014; Wiera and Mozrzymas, 2015). They also respond to neurodegenerative diseases, and are implicated in various psychiatric conditions (Pantazopoulos and Berretta, 2016; Pantazopoulos et al., 2015; Sorg et al., 2016; Testa et al., 2019; Wen et al., 2018).

The role of PNNs in the control of critical periods for plasticity is well established (Bernard and Prochiantz, 2016; Beurdeley et al., 2012; Lee et al., 2017; Liu et al., 2013; Mirzadeh et al., 2019; Miyata and Kitagawa, 2016), and PNN attenuation in various ways can restore plasticity in the adult CNS and enable recovery of function after injury (Carulli et al., 2010; Day et al., 2020; Duncan et al., 2019; Fawcett et al., 2019; Lensjo et al., 2017b; Romberg et al., 2013; Rosenzweig et al., 2019; Rowlands et al., 2018; Soleman et al., 2013; Yang et al., 2015). The effect of PNNs on plasticity depends not only on the various chondroitin sulphate proteoglycans (CSPGs), but particularly on their sulphated glycosaminoglycan chains (CS-GAGs). Chondroitinase ABC (ChABC) digests these CS-GAGs in PNNs and in the surrounding ECM giving a window of increased synaptic plasticity (Chu et al., 2018; Romberg et al., 2013; Yang et al., 2015). Absence of PNN components such as the Haplns that links CSPGS to hyaluronan, aggrecan or tenascin-R lead to attenuation or absence of PNNs, which affects critical periods and extends juvenile levels of plasticity into adulthood (Carulli et al., 2010; Giamanco et al., 2010; Gottschling et al., 2019; Morawski et al., 2014; Rowlands et al., 2018). Aggrecan is a CSPG that is abundant in PNNs and is a key activity-dependent component for formation of PNNs (Giamanco et al., 2010; Hou et al., 2017; Rowlands et al., 2018). Ablation of aggrecan prevents PNN formation, and in the visual cortex has shown to reinstate juvenile ocular dominance plasticity (Rowlands et al., 2018).

Recently, a role for PNNs and CSPGs in memory has emerged. In fear memory, which depends on the amygdala, local injection of ChABC enables erasure of fear memory in adult animals which is normally possible only in juveniles (Gogolla et al., 2009). Similarly, CSPG digestion has enabled extinction training of drug craving and seeking (Blacktop and Sorg, 2019; Blacktop et al., 2017; Slaker et al., 2015; Slaker et al., 2018; Xue et al., 2014). Object recognition memory is dependent on the perirhinal cortex, and here ChABC digestion prolongs memory retention, may enhance memory acquisition, and affects synaptic plasticity (Romberg et al., 2013; Yang et al., 2015).

PNNs have previously been implicated in long term memory (Tsien, 2013). Since memory is encoded in the pattern and strength of synaptic connections, and PNNs regulate synapse dynamics and plasticity, it has been proposed that the lattice-like structure of PNNs might encode long-term memory (Tsien, 2013). An attraction of this idea is that PNNs are relatively stable over time, and their persistence does not require energy. Synaptic withdrawal occurs under various circumstances including stroke, anoxia and hibernation. An unsolved question is how memories can remain stable after prolonged periods of brain anoxia, cooling or hibernation which cause extensive synaptic withdrawal (Arendt and Bullmann, 2013; le Feber et al., 2017; Sandvig et al., 2018; Xerri et al., 2014). Most mechanisms for synaptic maintenance require energy, yet PNNs remain in place after anoxia. Could they provide a substrate to stabilize synapses and enable memory retention. The hypothesis that PNNs regulate long term memory makes two predictions. The first is that memories will be reflected in the lattice pattern of PNNs, reflecting the pattern of synaptic connections. The second is that the persistence of memories after hibernation or anoxia will depend on the presence of PNNs. Our study addresses this second prediction.

Animal models of hibernation can address the mechanisms of re-wiring. One model of short-term hibernation-like state (HLS) involve administration of 5’-AMP combined with cooling to activate the hypothalamic nuclei for induction of torpor (Carlin et al., 2018; Carlin et al., 2017; Carlin et al., 2016; Kawamura et al., 2019). Induction of torpor/HLS? by this method in wild type (WT) mice produces a reversible withdrawal of synapses on cooling and rewarming. Under these circumstances, around 30% of synapses in the CA1 hippocampal region are withdrawn. In a few hours, the animals passively rewarm, and synapse number returns to near the initial level. Several intracellular players have been suggested to mediate this recovery (Bastide et al., 2017; Peretti et al., 2015), but the rewiring mechanism is only partially understood.

The overall aim of the study was to test the hypothesis that PNNs are necessary for the maintenance of long term memories. For this we used the model of synapse withdrawal caused by HLS. The study focused on hippocampal spatial memory, assessed using the Morris water maze (MWM). The involvement of the hippocampus in spatial and other types of memory has been shown by hippocampal lesions, inactivation by local anaesthetic, transgenic and optogenetic methods and transmitter blockade. All these can affect acquisition and retention of spatial memory. The hippocampus also contains place cells which are central to spatial memory (Broadbent et al., 2006; Kentros et al., 1998; Morris et al., 1982; O’Keefe, 1979; Tanaka et al., 2018) and contains prominent PNNs. We first measured synapse withdrawal and reconnection caused by imposing HLS. This was done in two models that attenuate PNNs, enzymatic digestion of PNNs, and local knockout of aggrecan via AAV1-*hSynapsin-Cre* injection in floxP *Acan* mice. Animals were trained in spatial memory in the MWM, cooled, then memory retention and ability to re-learn were measured in the presence and absence of PNNs. We found that HLS caused synapse withdrawal, particularly on the PNN-bearing PV^+^ inhibitory interneurons. Synapses reconnected after HLS, and the numbers of inhibitory and excitatory regenerated synapses were increased in the absence of PNNs. HLS led to partial loss of the memory and impaired subsequent re-learning. The absence of PNNs did not lead to a greater memory deficit compared to controls. Instead, absence of PNNs accelerated re-learning. We conclude that PNNs in the hippocampus are not required for the retention of spatial memories when the brain suffers an event that causes synaptic withdrawal.

## Results

**Scheme 1.**
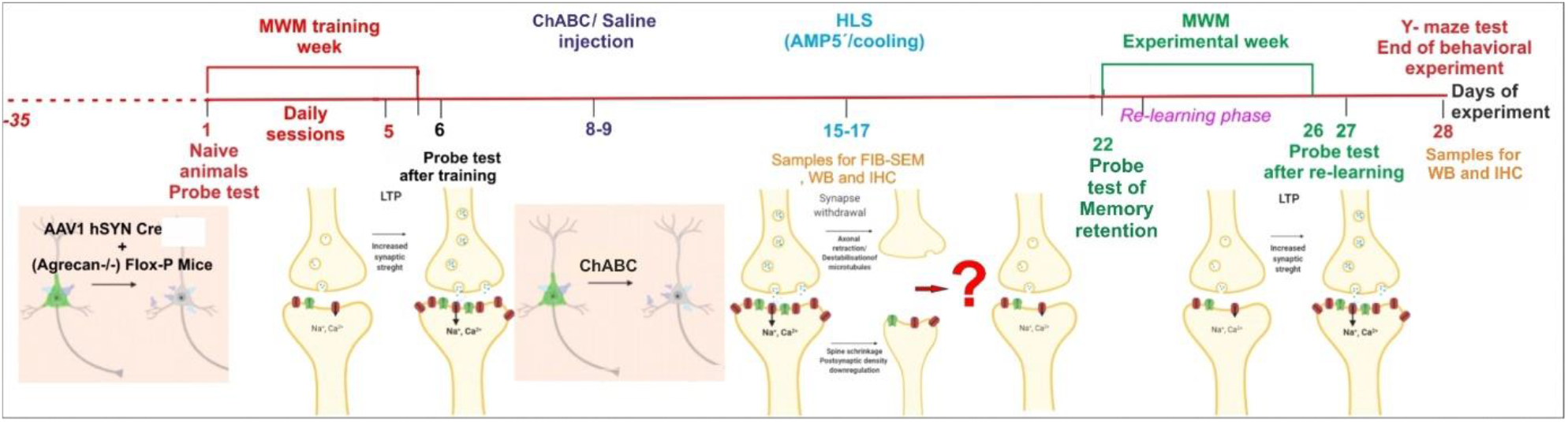
shows the chronological sequence of analyses used in the study.

### Study design and timeline

The aim of the study was to use models of spatial memory to determine whether the presence of PNNs in the hippocampus is required for restoration of memory after synapse withdrawal and replacement. Two methods of attenuating PNNs were used in two separate experiments: in the enzymatic ChABC experiment, ChABC was injected to 4 hippocampal sites bilaterally to digest PNNs, in the transgenic experiment conditional aggrecan knockout (AcanKO) animals received injections of AAV1-*hSynapsin-Cre* to both hippocampi five weeks before initial training (Scheme 1), eliminating PNNs (Figure S1; Injection sites at the end of the study, Figure S2; CA1 and PV^+^ neuron detail of WFA, PV^+^ signal). All animals were initially tested for MWM probe test to establish the level of the memory of the naïve animals, then they were trained for 5 days to learn the location of a submerged platform in the MWM. In the enzymatic experiment, ChABC or saline was injected on day 8-9, (ie 3 days after the training period). Half of the animals were placed in a HLS on day 15-17 to induce synaptic withdrawal. After one week to allow sufficient recovery, on day 22 a MWM probe test was done to analyse memory retention after HLS. The MWM was performed and repeated daily for 5 days in order to measure relearning. A probe test in which the submerged platform was removed was performed again at the end of this week on day 27. As a general test of hippocampal function, spontaneous alternation in the Y-maze was performed on day 28 after which the animals were perfused for histology. Statistical significance was marked ^#^p≈ 0.05, * p<0.05, **p<0.01, ***p<0.001 (for statistic, see table 1 in attachments).

### HLS temporarily reduces synapse numbers

The first step was to confirm that the induction of HLS leads to temporary synaptic withdrawal in the hippocampus as previously shown and to examine the effects on memory. Animals were subjected to a brief HLS, cooling to a core temperature of interval of 16-18°C, maintained for 45 minutes then re-warmed over 24 hours, as in the previous study (Peretti et al., 2015). The effect of the HLS on synapse numbers in the CA1 dendritic tree area was measured. Focused ion beam-scanning electron microscopy was used to measure overall synapse numbers. Sections of CA1 were taken from animals before HLS (37°C), immediately after HLS (16-18°C) and 24h after HLS by which time the body temperature had recovered to 37°C. In the sections, all synapses in selected areas were counted, defined by the co-location of a vesicle-containing presynaptic element and a thick or thin postsynaptic density on a dendritic figure. We observed that numbers of synapses decreased at the end of HLS, followed by restoration of synapse numbers at 24hrs to a slightly higher number (Figure 1A).

**Figure 1.**
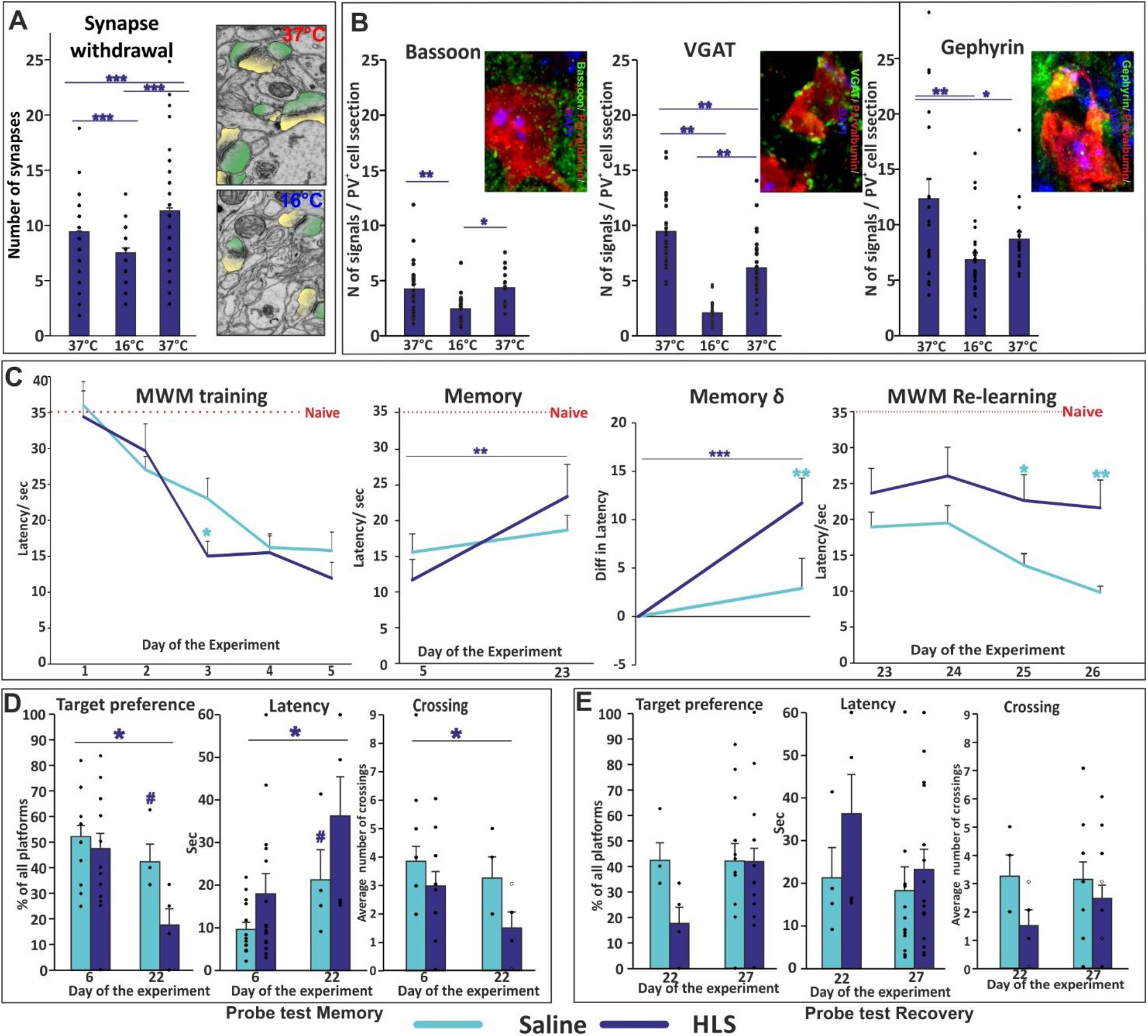
The synapse withdrawal effect of HLS was observed at CA1 level of hippocampus (A, representative image showing impact of HLS, pre- and post-synaptic site marked by yellow and green, respectively). Detail analysis of HLS on synapses targeting PV^+^ inhibitory interneurons body, showed significant withdrawal followed by complete recovery in Bassoon^+^ terminals, but only partial recovery at inhibitory synapses (vGAT and gephyrin) (B). Before the HLS procedure WT animals trained in Morris water maze (MWM) long-term memory task display normal learning curve. The memory retention was negatively affected by HLS and animals suffered with significant memory deficit, which was, nevertheless, not to the level of naïve animal performance. Additionally, animals after HLS have shown deficit in relearning of the MWM task, compared to WT animals (C). In the probe test trial, HLS animals showed significant loss of target preference, increased latency to reach the target and decreased frequency of crossing of target area. Additionally, strong trend toward significance in comparison to control group was observed in target preference and latency analysis (D). Probe test at the end of the study revealed the HLS animals have reached almost complete recovery in latency to reach the target zone and number of crossings and showed the same preference to target zone as the control animals (E). Statistical significance was marked #p≈0.05, *p<0.05, **p<0.01, ***p<0.001 (for statistic, see table 1 in attachments).

PNNs in the brain predominantly surround PV^+^ inhibitory interneurons. We performed analysis of individual pre and postsynaptic markers for excitatory and inhibitory synapses located on PV^+^ neurons, using bassoon as a presynaptic marker for excitatory synapses, and postsynaptic gephyrin and presynaptic vGAT to identify inhibitory synapses. The number of synaptic compartments on single optical sections of PV^+^ neuronal bodies was counted (Figure 1B). Just after HLS (16-18°C) a large reduction in bassoon, vGAT and gephyrin synapse number was detected (Figure 1B). After the 24hr passive rewarming phase (37°C) bassoon^+^ terminal numbers returned to normal and numbers of inhibitory synapses recovered to values slightly below pre-HLS. HLS also led to temporary reductions in several synapse-related proteins (see later).

#### HLS affects MWM memory

In order to study the effects of cooling and PNN manipulation and on long-term memory we tested place memory using the MWM. In this task animals learn the position of a refuge platform placed under the opaque water surface. This task is widely used to reveal spatial memory dependent on the hippocampus, and memory is adversely affected by various hippocampal interventions. After learning the position of the platform, the memory is usually stable for 4-5 weeks in normal animals.

The effect of HLS on memory in normal animals was assessed. Animals were trained to remember the position of the refuge platform by daily training for 5 consecutive days in the MWM, during which the latency time needed to find the target (first crossing of the platform boundary, with at least 0.5s spent at the target platform) was measured. Mice were then placed in an HLS in which their body temperature dropped to 16-18°C for 45 minutes, causing synaptic withdrawal as described above. Animals needed one week to recover from this procedure, after which their memory of the position of the platform in the MWM was tested again. For five further days animals were tested in the MWM and their ability to re-learn the refuge position was measured (Figure 1C). Probe tests in the absence of the platform were performed on the day 6 before HLS, 7 days after HLS and on the final day (Timeline Scheme 1, Figure 1D, E). During the 5 training days, animals showed a progressive decrease in latency for finding the target. Comparing memory before and after HLS, animals showed a significant increase in latency, although not to the pre-training level (Figure 1C). Memory loss was confirmed by comparing pre- and post-HLS probe tests, with changes in target preference, latency and target boundary crossing (Figure 1D). In the 5 days after the HLS period, HLS animals showed no significant shortening of latency while animals not subject to HLS continued to learn (Figure 1E). Overall, these results show that animals subjected to HLS demonstrated a period of synaptic withdrawal including withdrawal of synapses on PV^+^ interneurons, and a partial loss of MWM memory. After the HLS animals failed to learn, unlike non-cooled animals which continued to learn. Together these results show that HLS causes synaptic withdrawal followed by regeneration, and a partial memory deficit followed by impaired relearning.

### Removal of PNNs affects recovery of synapse numbers after HLS

The effect of PNNs on synapse numbers after recovery from HLS was examined. The ChABC group received injections to CA1 to digest the PNNs 1 week before HLS. PNN digestion lasts for over 3 weeks (Lin et al., 2008). Overall synapse number was measured by FIB-SEM. Both ChABC and control saline-treated animal groups had equally decreased numbers of synapses at the end of HLS, followed by restoration of synapse numbers at 24 hrs. ChABC injection influenced synaptic numbers. Before cooling there were more active synapses per section in the ChABC pre-treated group than in saline controls (Figure 2A). After rewarming synapse numbers increased in all groups. In the saline-treated group the synapse number was slightly higher than in the ChABC group.

**Figure 2.**
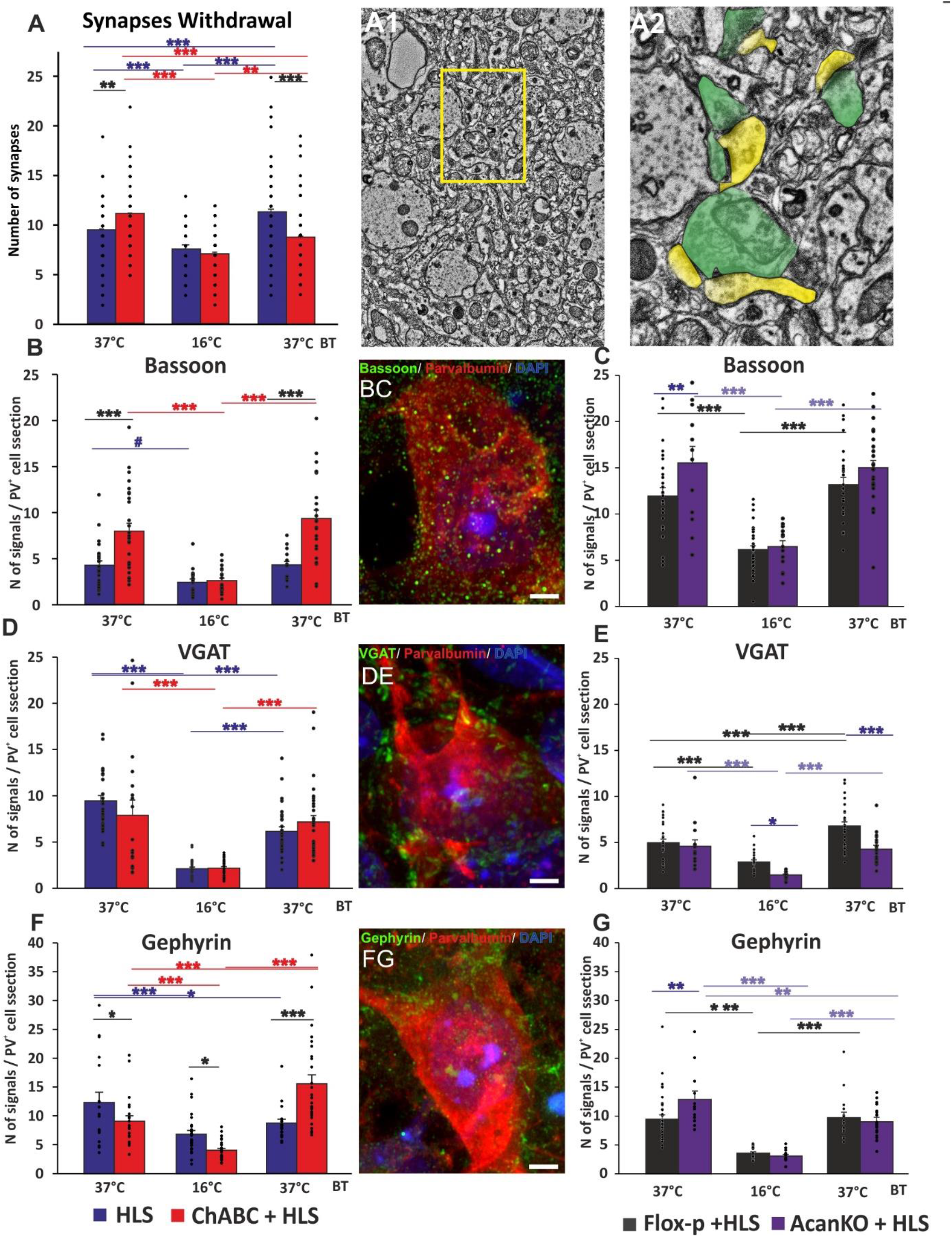
Focused ion beam-scanning electron microscopy (FIB-SEM) was used to investigate number of synapses (A1; detectable pre- and postsynaptic element -yellow and green respectively in A), PSD and ± vesicles) in the CA1 dendritic tree area before, immediately and 24hrs after HLS and to observe the impact of enzymatically digested PNNs. At the initial euthermic condition, the ChABC treated CA1 area showed increased number of synapses. Immediately after HLS in both groups significant drop in number of synapses was observed. After 24hr rewarming period both groups have recovered synapses to euthermic level, with ChABC group reaching only WT euthermic values (A, Illustration A1, detail A2). Furthermore, to investigate direct impact of HLS on parvalbumin^+^ inhibitory interneurons and role of intact PNNs in this process, confocal microscopy was used. In detail, average number of presynaptic terminals (bassoon (B, C)/vGAT (D, E)) and postsynaptic densities (gephyrin (F, G)) during euthermic conditions, immediately and 24h after cooling was measured. For analysis were taken only signals on cell body to avoid variance in the number and length of axons and dendrites. Like in FIB-SEM analysis the reduction of number of signals was observed in pre- and post-synaptic structures immediately after HLS and significant rate of recovery was observed 24h after. The impact of the PNN integrity was significant, with robust increase of excitatory terminals on PV neurones in comparison with intact PNNs and with tendency of decreased inhibitory terminals. Illustrative images of measured samples and discrimination of synaptic compartments are shown. Illustrative images bassoon/ gephyrin/ vGAT (green), parvalbumin (red), DAPI (blue). Scale bar 10μm. Statistical significance was marked * p<0.05, **p<0.01, ***p<0.001 (for statistic, see table 2 in attachments).

Synapse numbers on PV^+^ interneurons were also affected by ChABC digestion. Just after HLS (16-18°C) a large reduction in bassoon, vGAT and gephyrin synapse number was detected in ChABC and control groups. After the 24hr rewarming phase bassoon^+^ terminal numbers returned to normal in the saline control group, but in ChABC-treated animals the number of bassoon^+^ terminals was more than double that of the control group (p<0.001) (Figure 2B). The number of vGAT and gephyrin structures decreased considerably immediately after cooling. After 24hrs numbers of inhibitory synapses recovered to values slightly below pre-HLS levels in control, but gephyrin numbers were higher in ChABC-treated animals than in saline controls.

Synapse numbers on PV^+^ interneurons were also measured in animals in which PNNs were ablated by injection of AAV1-*hSynapsin-Cre* virus into the hippocampus of aggrecan (*Acan*-floxP ^+/+^) transgenic mice. After virus injection, the AcanKO and control animals showed an immediate decrease in the number of bassoon^+^ terminals after HLS and returned to its initial value after passive rewarming. Prior to HLS, brains lacking aggrecan had a higher number of bassoon^+^ terminals compared to floxP brains and this increase was sustained after recovery as a trend (Figure 2C). Animals lacking PNNs had reduced number of vGAT^+^ terminals, during and after the HLS (Figure 2E). The postsynaptic inhibitory marker gephyrin was increased relative to floxP control before HLS in the AcanKO group but there was no difference after recovery (Figure 2G).

At the end of the behavioural study 3 weeks after PNN digestion (day 28, 13 days after HLS) the effect of PNN attenuation persisted, with increased numbers of bassoon and gephyrin stained stained structures. (Figure 3, Figure S2). Together these results show that PNNs affect the recovery of synapse numbers after HLS, and this effect was much more marked on PV^+^ interneurons.

**Figure 3.**
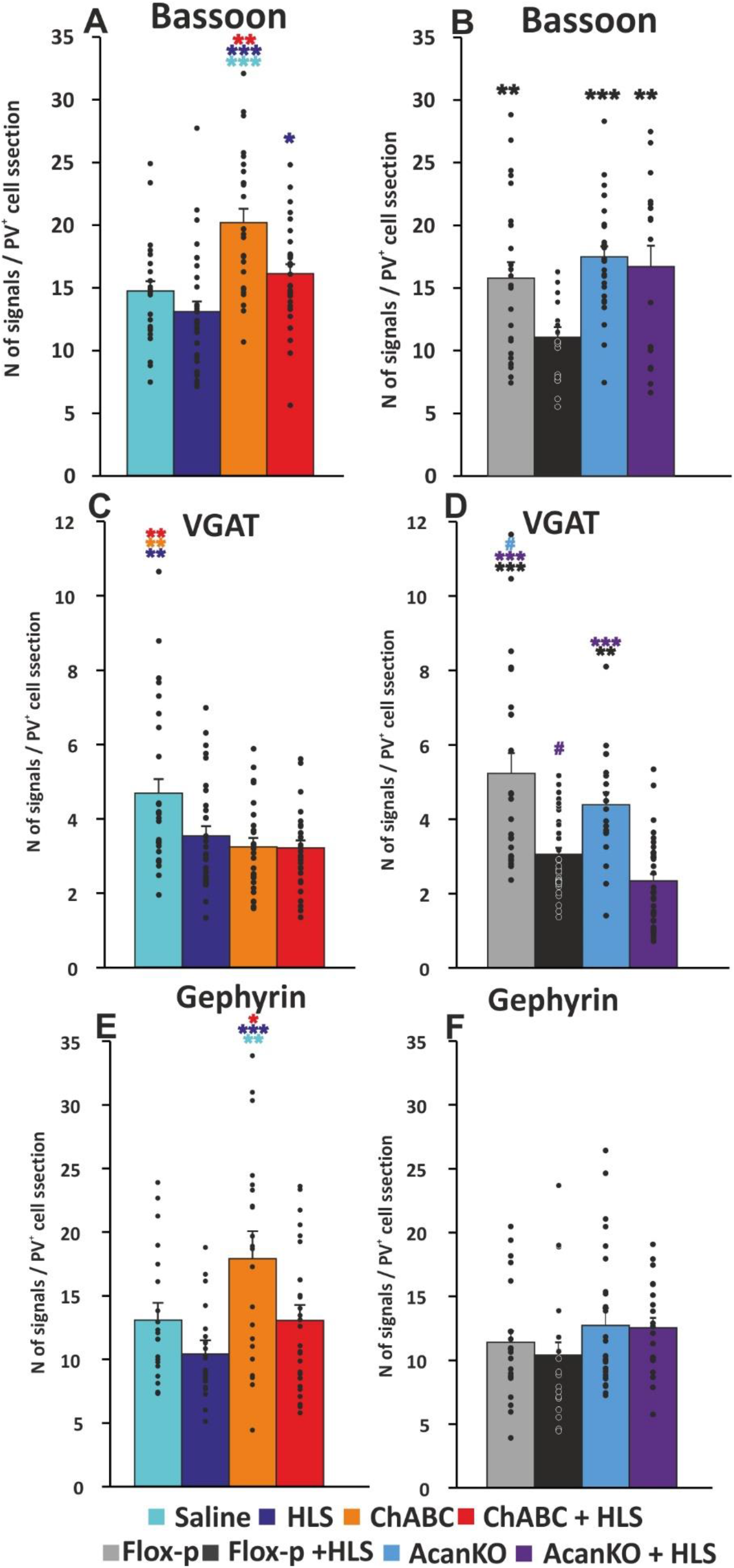
Number of excitatory and inhibitory inputs (all on CA1 PV^+^ neurons) was measured on day 28. Animals with digested PNNs have shown significantly higher number of bassoon^+^ pre-synaptic elements, when compared to other treatment groups. Moreover, the HLS animals pre-treated with ChABC has shown significantly prevented reduction in comparison with HLS only (**A**). All treatments, HLS, ChABC and their combination showed decreased amount of vGAT^+^ pre-synaptic elements in comparison with saline treated animals (**C**). On postsynaptic site the gephyrin molecules were significantly increased in animals from ChABC group in comparison with all other treatments. Minor reduction of gephyrin positivity was detected in HLS group (**E**). In the transgenic animals, strong effect of HLS was observed. The HLS treated floxP animals had significantly lower number of bassoon (B) than all the other groups, even the AcanKO + HLS. Additionally, the HLS condition led to and significant reduction of vGAT positive presynaptic elements, when compared to non-cooled control groups. Both AcanKO groups had lower amount of vGAT^+^ elements than related floxP groups and cooling further reduced these numbers (**D**). In gephyrin positive postsynaptic elements, contrary to ChABC study, no difference was observed between any of the transgenic animal group (**F**). Statistical significance was marked ^#^p≈ 0.05, * p<0.05, **p<0.01, ***p<0.001 (for statistic, see table 3 in attachments).

### Cleavage of PNNs affects synaptic protein levels after HLS

Synaptic protein changes after HLS were quantified by western blot analysis of pre- and post-synaptic markers (Figure 4). SNAP25 is presynaptic, vGLUT is in excitatory terminals, vGAT is in inhibitory teminals, GAD65/67 is present in inhibitory presynapses and cell bodies, PSD95 is postsynaptic. The proteins were analyzed in hippocampal tissue before HLS, during HLS and after 24hrs of re-warming. The level of all the markers except GAD65/67 decreased immediately after cooling, then vGLUT, SNAP25 and vGAT levels returned to normal with re-warming. PSD95 remained low after rewarming, while GAD65/67 increased slightly. ChABC pre-treatment led to increased SNAP25 on re-warming, and lower GAD65/67 compared to saline controls. This is consistent with increase vesicle secretion and lowered GABA inhibition.

**Figure 4.**
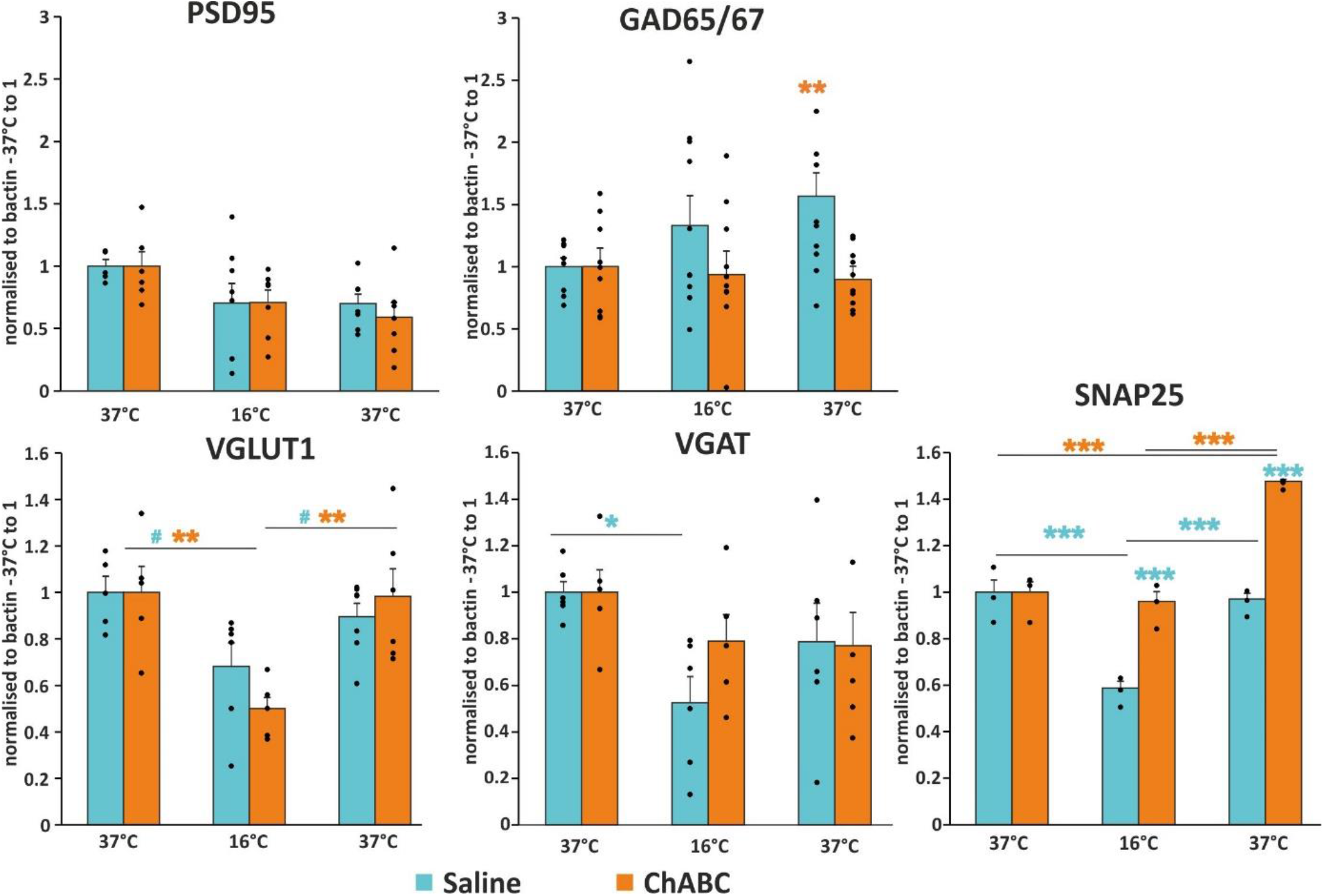
The western blot analysis of pre- and postsynaptic markers in euthermic animals immediately and 24h after HLS has showed similar shift observed in FIB-SEM and confocal microscopy. The whole hippocampal lysate is, however, less specific taking in account not only membrane bound proteins. In GAD65/67 the 24hr increase was observed in control animals whereas the SNAP25 as more general synaptic marker was showing the accumulation of the protein in ChABC treated animals in comparison with saline controls. Statistical significance was marked ^#^p≈ 0.05, * p<0.05, **p<0.01, ***p<0.001 (for statistic, see table 4 in attachments).

Analysis of synaptic proteins was performed also at the end of the study (timeline Scheme1 Figure 5). Animals that underwent HLS showed decreased levels of the GABA-producing enzyme GAD65/67 when compared to saline controls. ChABC treatment in non-hibernated animals caused an increase in the presynaptic markers vGLUT1, vGAT and SNAP25. ChABC digestion in HLS animals increased SNAP25 relative to saline-HLS animals.

**Figure 5.**
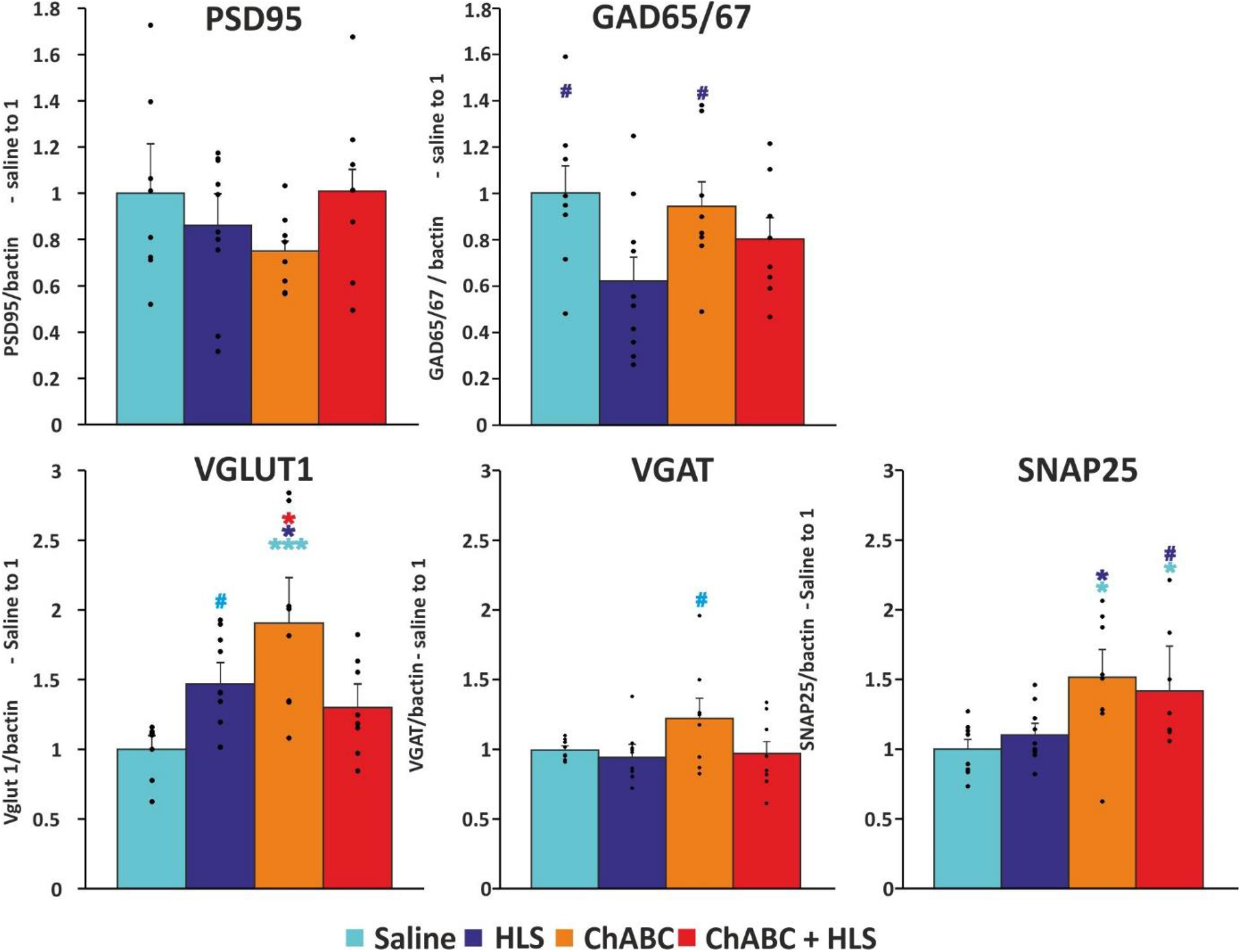
The western blot analysis of synaptic proteins in all experimental groups at the end of experiment has revealed trend in lover levels of postsynaptic proteins (PSD95, GAD65/67) after HLS partially compensated by ChABC pre-treatment. In presynaptic markers (vGLUT, vGAT and SNAP25) significantly higher protein levels were observed in ChABC treated animals; expression of SNAP25 marker was increased even after HLS state. Statistical significance was marked ^#^p≈ 0.05, * p<0.05, ***p<0.001 (for statistic, see Table 5 in attachments).

Overall, the results above confirm that subjecting animals to a hibernation-like state led to a temporary withdrawal of synapses in the hippocampus. Synapses were withdrawn from neurons in general and both excitatory and inhibitory synapses were withdrawn more markedly from PV^+^ interneurons. In the ensuing 24 hrs animals recovered and regained normal body temperature; during this time synapse numbers were partly or completely restored. Injection of ChABC into the hippocampus to digest PNNs had little effect on overall recovery of synapse number but a marked effect on synapses in PV^+^ interneurons, with an increase in both bassoon and gephyrin structures after rewarming compared to controls. Biochemical assay of synaptic proteins showed a similar pattern, with a decrease after HLS which was largely restored on rewarming. Prior ChABC treatment led to an increase after rewarming of SNAP25 and a decrease in the GABA-producing enzyme GAD65/67.

### PNN digestion or attenuation affects memory retention and re-learning

The results described above demonstrate that induction of HLS in mice leads to a partial loss of place memory after recovery, followed by a period of re-learning. We asked whether the presence of PNNs in the hippocampus would affect these processes. Two separate experiments were run, using different methods of PNN manipulation. 1) digestion of PNNs with chondroitinase ABC (ChABC) injected bilaterally into the hippocampus at day 8/9, just before HLS with saline injections as control (timeline Scheme1) (ChABC experiment) 2) local attenuation of PNNs in *Acan*-floxP animals through injection of AAV1-*hSynapsin-Cre* bilaterally into the hippocampus or control saline five weeks before the experiment (Acan experiment) (timeline Scheme1). In both these experiments removal of PNNs preceded HLS, but ChABC was injected after the initial learning phase just before HLS, while AAV1-*hSynapsin-Cre* was injected prior to initial learning. Thus, for the ChABC experiment all animals were untreated WT during the initial training period, but the aggrecan knockout animals had attenuated hippocampal PNNs throughout.

During the initial learning period the time taken to find the platform in the WT animals decreased from 33 s to 14 s. In the Acan experiment, AcanKO were injected with AAV1-*hSynapsin-Cre*, and floxP were saline-injected controls. There was no difference in their initial learning during the training phase, the time taken to find the platform decreased from 50 s to 24 s (Figure 6 A, E).

**Figure 6.**
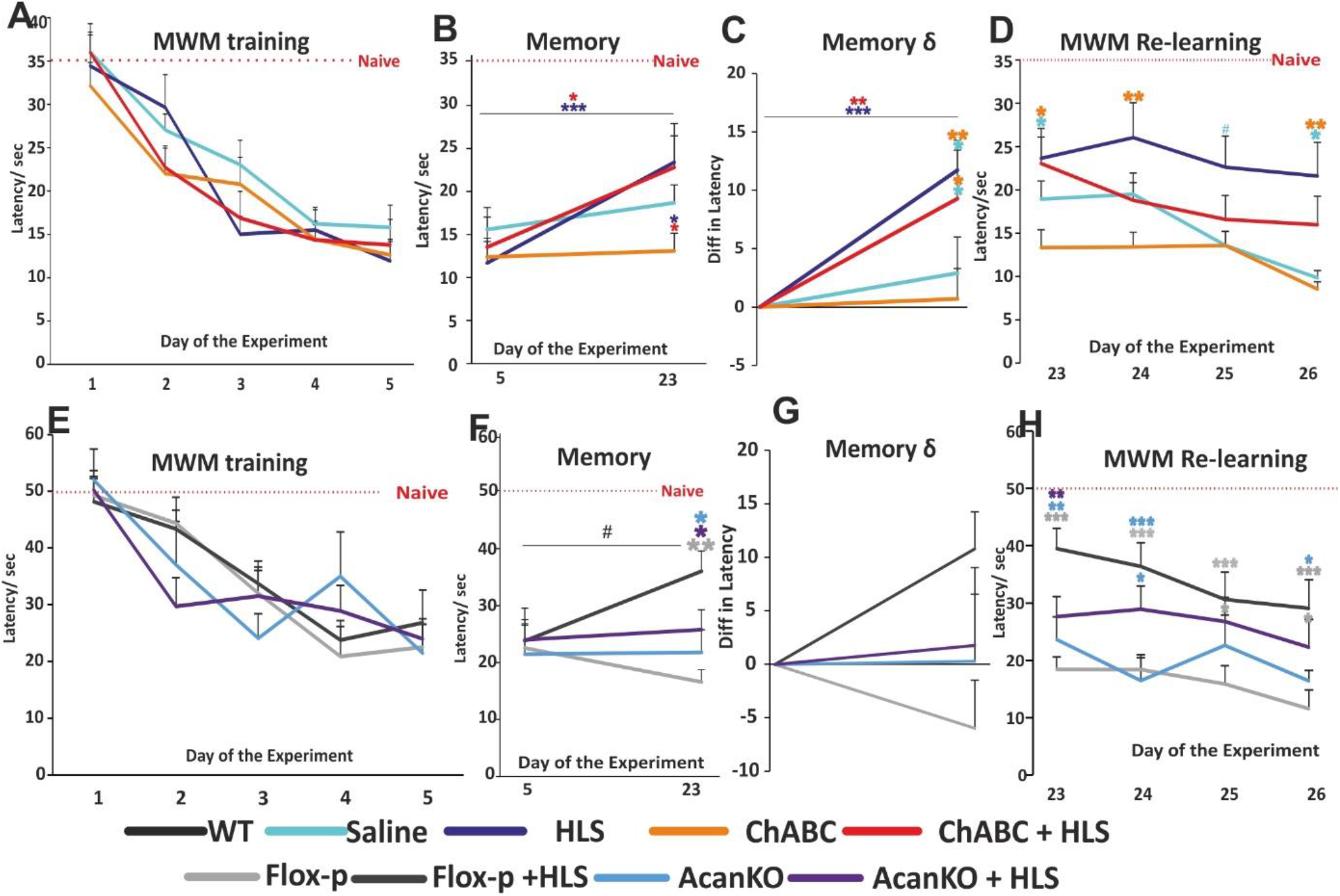
Morris Water maze test in long term memory setting has shown normal distribution and learning curve before the treatment application. All, WT, floxP and AcanKO mice were able to learn the MWM task (A, E). Training week was followed by week of ChABC injection (in WT group) and week of HLS. Applied procedures have led to partial loss of the memory, but not to the level of naïve animals (B, C). This loss was not observed in AcanKO + HLS mice (F, G). However, animals in HLS and AGG-KO + HLS group did not possess the ability to re-learn the task after deficit caused by cooling (D, H). Contrary, animals treated with ChABC showed to be fast learners despite the impact of HLS (D). floxP + HLS mice despite their memory deficit after cooling had ability to some extent re-learn the task, whereas both AcanKO and floxP animal groups showed lower memory deficit at the beginning of experimental week but their re-learning curve was either flat (AcanKO+HLS) or was not so pronounced as in case of ChABC treated animals (D, G). Statistical significance was marked ^#^p≈ 0.05, * p<0.05, **p<0.01, ***p<0.001 (for statistic, see table 6 in attachments).

#### Hibernation and PNN removal affect MWM memory

We next asked whether PNN attenuation affects the memory loss that occurs after HLS.

##### ChABC experiment

Half of the trained WT animals received ChABC injections into the hippocampus after the training period, the other half being injected with saline (ChABC n=29, saline n =29). The ChABC treatment caused loss of *Wisteria floribunda* agglutinin (WFA)-stained PNNs in the hippocampus (Figure S1A, in detail S2C, C1,2). After recovery from the surgery (1 week) half the ChABC and saline mice were placed in HLS, non-HLS animals serving as controls. Animals needed one week to recover from this procedure, after which their memory of the position of the platform in the MWM was tested again with a probe test in the absence of the platform. The two groups of animals that underwent HLS showed a partial loss of place memory, with no difference between ChABC and saline treated groups (Figure 6, B and C). Their latency time to find a target had increased, but not to the time score of naïve animals. However non-hibernated animals showed no deficit after the 2-week period (Figure 6, B and C). The probe test showed that HLS animals also had deficits in target preference, increased latency, and time spent in the correct quadrant (Figure 7).

**Figure 7.**
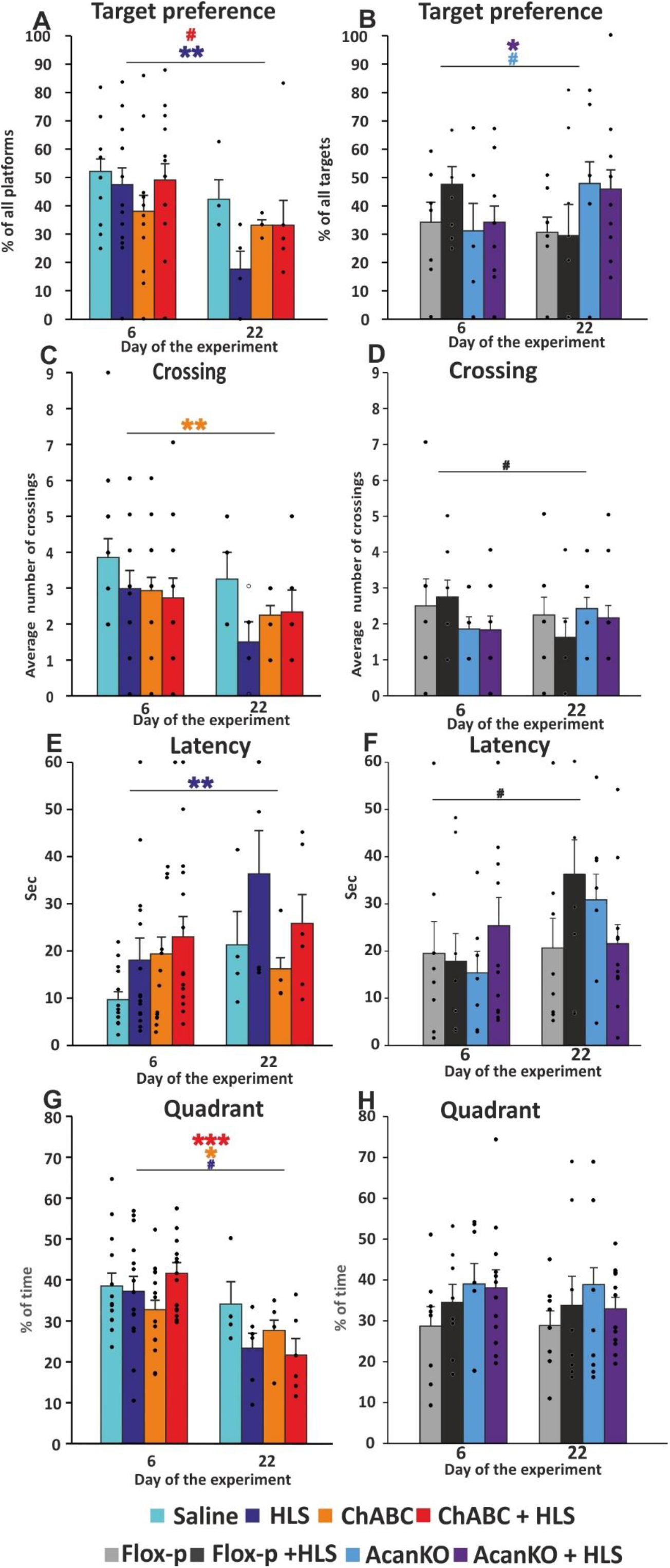
The probe test comparison of 6^th^ and 22^nd^ day of the study has showed the memory loss caused by HLS and preventive effect of either digestion of PNNs or the ablation of aggrecan production. Before the re-learning experiment, HLS animals had lesser target region preference than euthermic animals. ChABC treatment slightly mitigated the negative effect of hibernation (A). Increased preference of correct target zone was observed also in AcanKO ± HLS animals (B). Only ChABC treated and cooled floxP animals slightly decreased crossing of the target zone between test at day 6 and 22 (C, D).. The latency was significantly increased after HLS induction in both saline and floxP animals (E, F). Quadrant analysis has showed reduction in preference in ChABC ± HLS animals. Strong trend toward significance in saline HLS animals in reduction of quadrant preference was observed (G). Statistical significance was marked ^#^p≈ 0.05, * p<0.05, **p<0.01, ***p<0.001 (for statistic, see table 7 in attachments).

##### Acan experiment

One week after the last training session, half of the AcanKO (AcanKO n=19) and floxP (control, floxP n=16) animals underwent HLS the others serving as controls, then their memory in the MWM was tested one week later. Cooled floxP control animals showed a memory deficit, although not to the level of naïve animals. However hibernated AcanKO mice showed no deficit and their memory was at the same level as the last training day, suggesting that there had been rapid recovery in the week following HLS before testing. The two non-hibernated control groups also retained their spatial memory (Figure 6, F and G). The probe test showed that floxP control HLS animals had deficits in target preference, boundary crossing and latency, but the AcanKO animals had the same scores before and after HLS (Figure 7). These results show that HLS leads to a deficit in place memory although not to the level of naïve animals. AcanKO animals had recovered their memory in the week between HLS and the first MWM test.

### Absence of PNNs leads to accelerated re-learning after HLS memory deficit

The re-learning phase started one week after HLS, at which time the probe test described above was performed, followed with daily testing for a further 4 days to measure relearning at which point there was a further probe test (Scheme 1 timeline). In the ChABC experiment, the two groups that had not been subjected to HLS showed a further progressive shortening in their latency time to reach the target (Figure 6D). The two groups that received HLS started this period with a memory deficit compared to the end of the training period. Of these, the saline-treated controls showed no significant re-learning (Figure 6D). However, the ChABC pre-treated animals were able to re-learn the task, the slope of the learning curve being equivalent to that of the uncooled controls (Figure 6D). In the probe tests at the end of the study the ChABC groups (± HLS) showed improved target region crossings and latency to reach the target compared to the HLS saline controls. The unhibernated ChABC group scored above the saline controls on target preference, crossing and latency. (Figure 8). Effects of ChABC on PNNs last for at least 3 weeks due to persistence of the enzyme and slow PNN replacement (Lin et al., 2008), so the relearning enhancement occurs during PNN attenuation.

**Figure 8.**
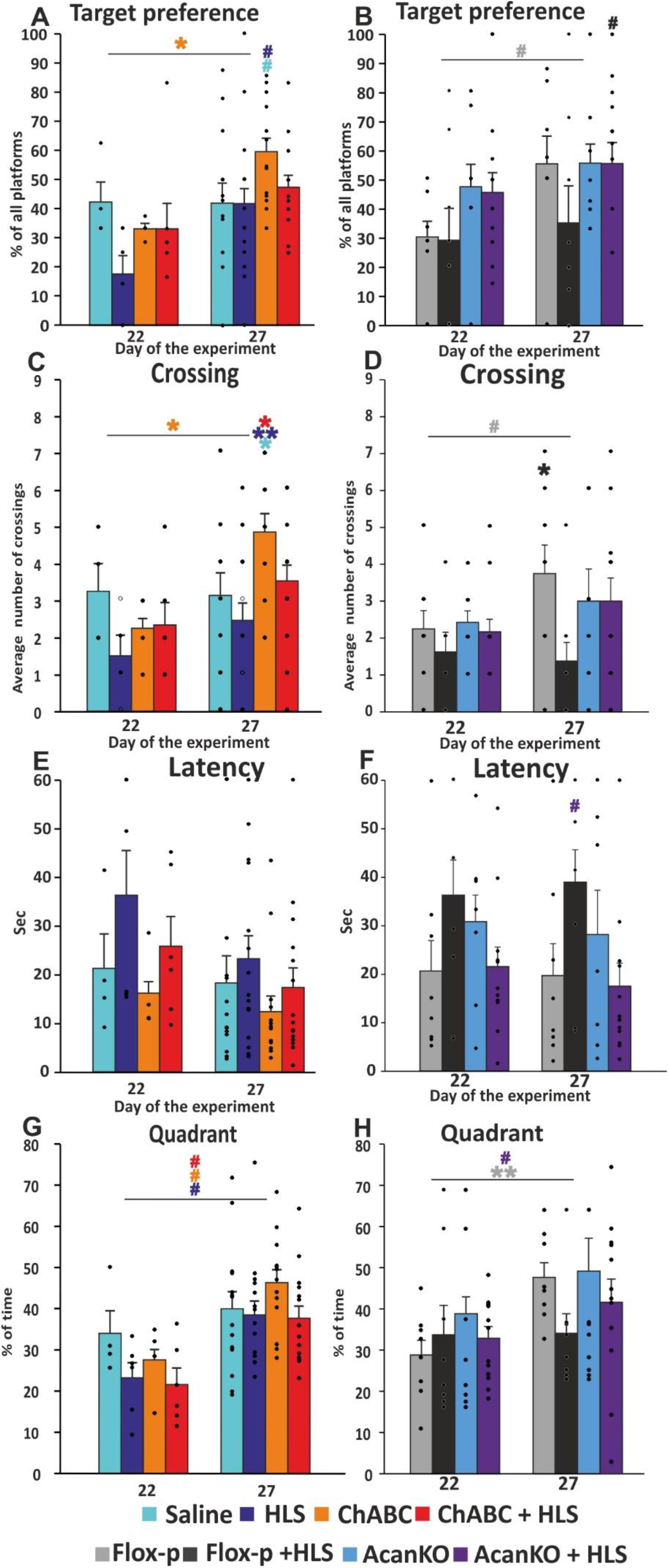
The relearning probe test comparison between 22^nd^ and 27^th^ day of the study. The ChABC treatment has shown to increase with strong trend target preference over the re-learning week, when compared with saline and HLS group at day27 (A). In AcanKO study, trend toward significance in target preference improvement was observed in floxP animals. Additionally, at the end of study the HLS state did not affect AcanKO mice when compared to floxP controls (B). In the number of crossings of target zone, the major improvement was observed in ChABC group. At 27^th^ day statistically higher frequency of crossings was observed, when compared to other treatment groups (C). In AcanKO study significantly more crossings were detected in euthermic floxP animals when compared to their cooled littermates at the 27^th^ day analysis (D). Animals with enzymatically or genetically removed PNNs had shorter latency to reach the target even at the 27^th^ time point, with strong trend toward significance in AcanKO group (E, F). In quadrant analysis the only significant improvement was detected in floxP control animals (H). Statistical significance was marked ^#^p≈ 0.05, * p<0.05, **p<0.01, ***p<0.001 (for statistic, see table 8 in attachments).

In the AcanKO experiment, the floxP + HLS controls started the re-learning period with a memory deficit, after which all the animals showed similar learning curve slopes. The AcanKO + HLS animals started this period with no deficit, suggesting that they had already compensated for the effect of HLS before the first post-HLS test, and then continued to improve (Figure 6F and H, Figure 8). In the probe tests at the end of the re-learning week, the floxP +HLS group performed consistently worse than the other groups, and the AcanKO + HLS group performed similarly to the unhibernated animals (Figure 6H, Figure 8).

Together, these results show that HLS produces a memory deficit although not a complete loss of memory to the level of naïve animals. HLS also adversely affects subsequent re-learning. In the two groups of animals that lacked PNNs due to ChABC treatment or transgenic deletion of Acan the ability of animals to recover their memory and relearn after HLS was enhanced so that these animals eventually performed similarly to the non-HLS groups. The effect on relearning is consistent with previous studies that show enhanced learning and memory retention following PNN attenuation (Fawcett et al., 2019).

An alternative test of ventral hippocampal memory function is spontaneous alternation, in which animals are placed in a Y-maze and their spontaneous entry in the arms is recorded. A hippocampal deficit can cause animals to re-enter the same arm rather than alternating between arms. This test was performed at the end of the re-learning week. We did not find any difference between any of the experimental groups (ChABC or AcanKO experiment, Figure S2). This indicates that by one week after hibernation hippocampal function is normal.

## Discussion

PNNs form a lattice around PV^+^ interneurons and some other neurons. Because PNNs contain inhibitory CSPGs, it is probable that regenerated synapses will tend to reconnect to the areas of naked neuronal membrane that they previously occupied rather than penetrating the PNNs to terminate elsewhere. This led to the idea that PNNs might specify memory by constraining the positions of synapses, and might therefore be the substrate for stable memories following conditions that lead to synapse withdrawal. Our experiments were designed to test this hypothesis for hippocampal-based place memory. The first step was to confirm that HLS leads to synapse withdrawal followed by reconnection in our model, as previously shown (Peretti et al., 2015). HLS led to synaptic withdrawal in the hippocampus, particularly on PV^+^ inhibitory interneurons, followed by restoration to near the original level. Removal of PNNs led to increased numbers of bassoon^+^ excitatory and gephyrin^+^ inhibitory synapses on PV^+^ interneurons following the regeneration phase 24hrs after hibernation. We then asked whether this HLS-induced synaptic withdrawal would lead to memory loss. We recorded partial loss of MWM place memory after HLS, although not to the level of naïve animals. Step 3 was to ask whether removal of PNNs before HLS would lead to a greater loss of MWM place memory than in animals with intact PNNs after recovery and synaptic reconnection. This did not happen. Digestion or deletion of ECM/PNNs did not increase the memory loss. In the ensuing four days animals were run daily in the MWM. HLS animals showed little re-learning during this time, but HLS animals with digested or knockdown PNNs relearned platform position at a normal rate. The fastest re-learning was in AcanKO animals lacking PNNs which recovered memory during the week between HLS and the post-HLS probe test. The overall conclusion is that HLS causes synapse withdrawal and partial loss of place memory; removal of PNNs in the hippocampus did not worsen this memory loss, although it increased the total number of synapses and enhanced subsequent re-learning. PNN removal therefore increases learning, as in previous studies (Fawcett et al., 2019).

Several PNN modification experiments have focused on the hippocampus. Animals lacking tenascin-R have attenuated PNNs, and these animals showed normal learning rates but increased agility in learning new platform positions (Morellini et al., 2010). On the contrary, animals with increased numbers and thickness of PNNs showed slow place learning, normalized after ChABC digestion in the hippocampus (Bertocchi et al., 2020). PNN digestion also affects hippocampal physiology; digestion of PNNs in CA1 restored long term depolarisation (LTD) that is lost in aged mice and enhanced excitability (Khoo et al., 2019), and both ChABC and tenascin-R knockout decrease CA1 long term potentiation (LTP) (Bukalo et al., 2001) In CA2, LTP and excitability were increased by digestion of PNNs, which in this region surround pyramidal cells (Carstens et al., 2016; Hayani et al., 2018).

The studies above demonstrate that PNNs in the hippocampus are involved in various memory functions. However, they do not clarify the involvement of PNNs in defining long-term memory, and particularly whether they can act as a persistent memory backup following events that lead to a loss of metabolism to power active synaptic maintenance. In our study, the animals lacking PNNs did not show a greater memory loss after HLS, indicating that hippocampal PNNs are not the substrate for stable place memory. All animals learned platform position over 5 days in the MWM, and lack of PNNs due to ChABC or AcanKO did not affect this process, as in tenascin-R knockouts (Morellini et al., 2010). Half the animals then underwent HLS, preceded by ChABC injection in the chondroitinase experiment. After 6 days recovery animals were again tested in the MWM to see whether they still remembered the position of the platform. Place memory is stable for months, so euthermic animals showed the same MWM results as before the time gap for operations. However, three of the HLS groups showed a clear loss of memory, shown in measures of target preference, boundary crossing, latency and quadrant time. In the AcanKO experiment, knockout HLS animals performed at the same level as euthermic controls, suggesting that they had rapidly recovered from the synapse withdrawal deficit during the week between HLS and testing. During the following 4 days animals were assessed for their ability to relearn, receiving daily MWM testing followed at the end by another probe test. Comparing day 22 and 27 probe tests, HLS animals with normal PNNs showed minimal relearning, particularly the *Acan-*floxP mice. However, the HLS animals lacking PNNs performed similarly to non-HLS controls. Importantly, the AcanKO + HLS animals recovered normal memory before the first post-HLS probe test. These relearning results match with previous work in which lack of PNNs has had a positive effect on memory acquisition and retention (Romberg et al., 2013; Rowlands et al., 2018; Yang et al., 2015).

Our experiments demonstrate that PNNs in the hippocampus are not necessary for retention of memory after synapse withdrawal. The two ways in which we depleted PNNs have different targets. ChABC digests all CS-GAGs, affecting the condensed PNNs visible with WFA staining, but also all the CS-GAGs in the more diffuse matrix that surrounds all synapses. Aggrecan knockout affects the condensed PNNs, for which aggrecan is a necessary structural component (Morawski et al., 2012; Rowlands et al., 2018). The results of both PNN-attenuation interventions were similar, with enhanced relearning from the memory deficit caused by HLS and enhanced synapse numbers on PV^+^ interneurons. This result excludes hippocampal PNNs and CS-GAGs from a role as backup memory storage devices. While the hippocampus has been accepted for many years as the central site for place memory, other sites input to the hippocampus and store place information. The entorhinal cortex contains grid cells which connect to hippocampal place neurons, and the cingulate cortex and some subcortical structures are also implicated (Christensen et al., 2020; Nelson et al., 2015; O’Mara and Aggleton, 2019; Steullet et al., 2014). An alternative hypothesis for the role of the hippocampus is as an indexer of distributed storage (Tanaka and McHugh, 2018). In our experiments, the AcanKO + HLS group regained their pre-HLS level of memory during the recovery week, without re-exposure to the MWM, suggesting that a memory backup inside or outside the hippocampus exists which is independent of hippocampal PNNs.

The short-term HLS model used in this study was developed to study the effects of neurodegenerative conditions on synapse regeneration (Peretti et al., 2015), and reproduces the synaptic withdrawal observed in hibernating mammals (Arendt and Bullmann, 2013; Carlin et al., 2018; Carlin et al., 2017; Carlin et al., 2016). The same withdrawal and reconnection of synapses was observed in the current study (Figure 1A). Recently, in the hibernating mammal the impact of seasonal rhythms and torpid state on the PNN intensity and distribution was examined (Marchand and Schwartz, 2020). No significant changes in memory were observed. In agreement with this study, we did not observe significant changes in PNN staining intensity in CA1 region of hippocampus, our region of interest, after HLS (Figure S2, F and G). The protection mechanisms of PNNs are well described in neurodegenerative diseases and oxidative stress and could be significant in hibernation and other conditions that cause synaptic withdrawal (Beurdeley et al., 2012; Hou et al., 2017; Reichelt et al., 2019).

In the previous study of synaptic withdrawal and regeneration with HLS, synapses in CA1 were counted by electron microscopy (Peretti et al., 2015). We replicated these counts, finding a decrease in overall synapse count and recovery 24 hours later as before. Pre-treatment with ChABC to digest PNNs and the general interstitial ECM did not affect overall synapse numbers although SNAP25 levels were increased (Figure 2A and 4). However, this analysis was neuron and synapse type unspecific, so we focused on PV^+^ interneurons the majority of which are surrounded by PNNs. We measured numbers of excitatory and inhibitory synapses on these cells, using bassoon, vGAT and gephyrin as markers, and investigating animals pre-treated with ChABC or AAV1-*hSynapsin-Cre* to cause aggrecan knockout (Figure 2, B-G). Both interventions affected synapse numbers prior to HLS, with increases in bassoon terminals. HLS caused a decrease in bassoon, vGAT and gephyrin of 45%. 78% and 45% respectively. 24 hrs later synapse numbers had largely recovered. Treatment with ChABC led to increased bassoon and gephyrin synapses on HLS recovery, although this was not seen in AcanKO (Figure 2, B and C) and this effect persisted for next two weeks with a lesser effect in the AcanKO group (Figure 3, A and B). PNN digestion or modification has previously shown a significant impact on the number of excitatory and inhibitory inputs on PV^+^ inhibitory interneurons (Donato et al., 2013; Gottschling et al., 2019), and to affect LTD, LTP and network activity in the cortex (Fyhn et al., 2004; Lensjo et al., 2017b; Romberg et al., 2013; Shi et al., 2019; Thompson et al., 2018). As with most studies, we have focused on the dense PNNs that surround PV^+^ interneurons and some other neurons, and which are visualized by WFA or aggrecan staining. However every synapse is embedded in CSPG-rich extracellular matrix, although not condensed through linkage of CSPGs, hyaluronan and Haplns (Fawcett et al., 2019; Kwok et al., 2014).

There have been various studies of the effect of PNN manipulation on memory acquisition and retention, but none in the context of acute synaptic withdrawal. The first study used fear conditioning as the model, with digestion of PNNs in the amygdala. Here, PNNs in adults make memories resistant to erasure by extinction training, while PNN digestion re-enabled the immature situation where fear memory can be erased, a form of reverse learning (Gogolla et al., 2009). Digestion of PNNs in the auditory system also led to increased learning flexibility (Happel et al., 2014). Digestion of PNNs in secondary visual cortex disrupted fear memory recall, probably by destabilizing network synchronization (Thompson et al., 2018), and removal of hippocampal PNNs disrupted contextual and trace fear memory (Hylin et al., 2013). PNNs in the hippocampus and cingulate cortex have been linked to fear memory consolidation, and PNN numbers actually increased in the hippocampus after fear conditioning, and increasing PNN numbers by transduction with Hapln-1 enhanced memory (Shi et al., 2019). Several studies have used novel object recognition (NOR), a task dependent on perirhinal cortex, as the memory model. Here, digestion of PNNs with ChABC or attenuation of PNNs through *Ctrl1* knockout, led to prolongation of object memory coupled with increased LTD in the perirhinal cortex (Romberg et al., 2013). Similarly, ChABC or treatment with an antibody recognising the inhibitory 4-sulfated CSPGs found in PNNs, was able to restore normal NOR memory in an Alzheimer models (P301S tauopathy and the APP/PS1 amyloid model).(Vegh et al., 2014; Yang et al., 2015; Yang et al., 2017). PNN digestion also has effects on addiction memory (Sorg et al., 2016)),

Overall, our experiments confirm that HLS leads to a temporary global withdrawal of synapses in the CA1 region of the hippocampus followed by regeneration. We show that synaptic withdrawal was particularly marked on PV+ interneurons most of which bear PNNs. Digestion of the PNNs enhanced numbers of excitatory and inhibitory synapses on these cells. Disruption of PNNs in the hippocampus did not affect MWM learning, and did not increase memory loss immediately after HLS. However there was a positive effect on recovery of memory and relearning in the days after HLS.

## Methods

### Animals

Experiments using enzymatic digestion (Chondroitinase ABC) of PNNs were conducted on males with C57Bl/6 background (age = 14 weeks, weight = 27±2 g, n=112). Animals for behavioral experiments were separated in following four groups; saline (n=14), hibernated like state (HLS, n=15), ChABC (n=14), ChABC + HLS (n=15), from which, additionally, IHC and WB analysis (Both n=4/group) was performed. Animals for FIB-SEM microscopy (n=2/group/ time point), acute IHC (n=3/group/ time point), acute WB (n=4/group/ time point) were used to observe the dynamic changes in synaptosome.

Experiments with local Acan knockout via AAV1-*hSynapsin-Cre* is achieved with stereotaxic hippocampal injections on *Acan* GT3^+/+^/GT5^+/+^ floxP mice with C57Bl/6 background (age=14 weeks, w=27±2 g, n= 53). Animals for behavioral experiments were separated in following four groups; saline floxP (n=8), floxP + HLS (n=8), AcanKO (n=7), AcanKO + HLS (n=12), from which, additionally, IHC and WB analysis (both n=4/group) was performed. Animals for acute IHC (n=3/group/ time point) were used to observe the dynamic changes in synaptosome.

All experiments were performed in accordance with the European Communities Council Directive of 22 September 2010 (2010/63/EU), regarding the use of animals in research and were approved by the Ethics Committee of the Institute of Experimental Medicine ASCR, Prague, Czech Republic.

### ChABC injection

One week after MWM training period (Scheme 1) animals received ChABC (Sigma-Aldrich, Germany) or saline injections in to both hippocampi. Semiautomatic motorized operating system with mice 3D brain atlas (Neurostar, Tubingen, Germany), was used to drill the skull and inject the treatment within the pre-estimated coordinates to fully cover the volume of hippocampus. Animals were under isoflurane anaesthesia, receiving local painkillers (mesocaine, subcutaneous 30 µl), shaved, and cut alongside sagittal suture to open 1cm length window into the skin. System was accustomed on brain marks (Bregma, Lambda) and skull rotation, in order to predict injection path. With low Z axes speed (1 mm/min) the driller prepared 4 scull micro inserts for each hippocampus. With low speed the injector was set in the coordinates. Injection parameters were estimated; volume = 0.6 μl/injection point, speed = 0.1 μl/min. After each injection the 3 min steady interval before taking out the injection needle was set, to prevent leaking. The skin was sutured and treated to prevent reopening (Novikov, Prague Czech Republic).

### Induction of local Acan knockout

Five weeks before MWM training *Acan* floxP animals received stereotaxic injections of AAV1-*hSynapsin-Cre* virus (c= 1_x_10^13^ u, volume = 0.6 μl/injection point, speed =0.1 μl/min / saline control) to knockdown production of Aggrecan and destabilize PNNs in both hippocampi. The same operation parameters and stereotaxic coordinates were used as in case of ChABC treatment.

### AMP mediated Hibernation like state

One week after ChABC injection (or 7 weeks after Acan knockout induction), the mice underwent HLS protocol (see in detail; (Peretti et al., 2015)) Mice were intraperitoneally injected with 5′AMP (0.1 g/ml, 0.5 μl/g, Sigma-Aldrich) at room temperature. Body temperature was measured per rectum before injection and then in 15 min interval until the mice reached the room temperature (25°C ± 1). The breathing depth and frequency were controlled. After reaching the room temperature the mice were placed in the stable cold environment (4°C) and carefully watched. When the body temperature reached 18°C, the time interval of 45 min necessary to induce synapse retraction was measured. Later the mice were placed at room temperature to passively warm up.

### FIB-SEM microscopy

#### Sample fixation and resin embedding

For FIB-SEM microscopy the mice were perfused with sodium cacodylate buffer (pH 7.4) followed by perfusing solution (2.5% glutaraldehyde/ 2% paraformaldehyde in Na cacodylate buffer). The whole mouse brain was fixed for at least 5 hours in 2% paraformaldehyde and 2.5% glutaraldehyde in 0.1 M CDS (cacodylate buffer) at a pH of 7.4. The fixed tissue was cut with vibratome (Leica VT1200) into 150 µm thickness slices and they were placed in 10 ml glass vials with fresh solution of fixatives. Brain slices were fixed for 1 hour on ice. The samples were rinsed in 0.1M CDS three times, 5 minutes each. Then the samples were stained with 1.5% (w/v) potassium ferrocyanide and 1% (w/v) osmium tetroxide in 0.1 M CDS for 30 minutes and next with 1% (w/v) osmium tetroxide in 0.1 M CDS for 30 minutes. After postfixation and staining samples were rinsed in distilled water three times, 3 minutes each. Then the samples were stained with 1 % (w/v) uranyl acetate in distilled water for 30 minutes and rinsed in distilled water two times, 5 minutes each. Brain slices were dehydrated in graded ethanol (EtOH) series, 2 minutes each change (1x 30 %, 1 x 50 %, 1 x 70 %, 1 x 95 %, 2 x 100 %), in 100 % EtOH 5 minutes. The dehydration was followed by embedding in mixture of Epon EmBed 812 hard with EtOH (1:1) for 30 minutes and then in 90 % Epon EmBed 812 hard an overnight on rotator. Then we changed Epon with the fresh one (100 %) and agitated slowly for 4 hours. Finally, the sections were placed on glass microscopes slides coated with mould separating agent using wooden cocktail sticks and covered with fresh 100 % Epon and placed in 60° C oven for 24 hours.

#### Preparing the sample for the FIB/SEM

After polymerization, the resin layer containing the samples was separated from between the two glass microscope slides and washed thoroughly to remove any mould separating agent. Using a transmitted light microscope (Carl Zeiss Axiozoom.V16) we identified the region of interest on mouse brain slices within the slice of resin. A small (5 mm x 5 mm) square around the region of interest was cut using a razor blade and stuck to the top of blank resin block with acrylic glue. After the glue hardening the block was mounted into holder of the ultramicrotome (Leica EM UC7) and trimmed a small pyramid around the region of interest with a razor blade. Then we trimmed the block surface with glass knife until the embedded tissue will appeared on the resin surface. Finally, the trimmed block was cut away from the remaining resin stub to ensure that only a small block is placed inside the FIB-SEM. The small epon block was mounted on a regular SEM stubs using conductive carbon and coated with 25 nm of platinum (using High Vacuum Coater, Leica ACE600).

#### FIB-SEM imaging

Ion milling and image acquisition was performed in Dual beam system FEI Helios NanoLab 660 G3 UC.

By using a low magnification and secondary electron imaging (20 kV and 0.8 nA) the block was oriented into best position and the region of interest was chosen. The protective layer (approximately 1 um thick) of platinum was deposited, using the gas injection system of the microscope, onto the surface of the block, above the region of interest. A large trench around protective region was milled at a current of 21nA and 30kV by focus ion beam. It was followed by fine milling at 0,79 nA and 30 kV, thickness of slices was 90 nm. The SEM imaging of the milled face (area of interest) was done in backscattered imaging mode and the serial SEM images were acquired at 2 kV and 0.2 nA using an InColumn backscattered electron detector (ICD). The XY pixel size was set to 3 nm.

### Immunohistochemistry

#### Tissue isolation and preparation

For immunohistochemistry, the mice were transcardialy perfused with PBS followed by paraformaldehyde (4% PFA in PBS). The brains were left in PFA overnight then gently washout in PBS, 20% sucrose and then 30% sucrose (each solution time= 24 h, t=4°C). Using cryotome 20um thick coronal sections were prepared.

#### IHC Staining and visualization

To evaluate the effect of treatments on PNNs and synaptic input of parvalbumin positive inhibitory interneurons, immunohistochemical staining on series of 20μm coronal brain section was performed.

To observe integrity of PNNs and monitor the effect of ChABC or AAV1*hSynapsinCre* local knockout staining biotinylated *Wisteria floribunda* agglutinin (WFA, 1:150, Sigma, Germany), and anti-aggrecan primary antibody (1:150, Sigma) together with anti-parvalbumin primary antibody (1:500, Synaptic Systems, Goettigen, Germany) were used. For analysis of parvalbumin positive inhibitory interneurons synaptic connectivity, antibodies against parvalbumin (1:500, Synaptic Systems), gephyrin (1:250, inh. Postsynaptic, Synaptic Systems), bassoon (1:250, Excyt. presynaptic active zone, Synaptic Systems) and vGAT (1:200, inh. Presynaptic, Synaptic Systems) were used. To visualize primary antibodies positivity anti-mouse Alexa Fluor 594 (1:200, on PV neurons) and anti-rabbit Alexa Fluor 488 (1:300, on gephyrin, bassoon, vGAT) secondary antibodies were used. To visualize the primary antibody a streptavidin 488. Alexa Fluor 488 and alexa Fluor 594 secondary antibodies were used (goat anti mouse 1:300, goat anti rabbit 1:200, streptavidin AF488 1:300).

#### Confocal imaging

To evaluate synaptic markers, present on parvalbumin^+^ inhibitory interneurons in CA1 region of hippocampus, confocal microscope Zeiss880Airyscan was used. The 63x oil immersion objective (Plan-Apochromat 63x/1.4 Oil DIC M27, zoom 1.5, pixel dwell 0,82 μs, x: 1272, y: 1272, z: 58, channels: 3, 12-bit). To visualise intensity and structure of PNN signal by WFA and aggrecan antibody staining, 10x objective (plan-achromat10x/0.45 M27, zoom 0.7, Pixel dwell 0.77μs, 1214×1214×22, Average-line 2, 12-bit) and 60x oil immersion objective (Plan-Apochromat 63/1.4oil DIC M27, zoom 1.5, Pixel dwell 0.82, 1272×1272×56, Average-line 2, 12-bit) were used. Lasers-track 1 (488 nm, first detector), track 2 (561 nm) and track 3 (405 nm) (both second detector) with two detectors setting. Laser power was maintained below 3%.

### Western blot

#### Tissue isolation and preparation

For western blot analysis the tissue was prepared on ice within the tissue lysate solution (ddH_2_O, Tris 50 mM, NaCl 150 mM, EDTA 2.5 mM, Triton X 1%, sodium deoxycholate 0,1%, PhosSTOP and Complete-mini EDTA-free (Roche, Germany)). The brain was cut to hemispheres and the hippocampus isolated and placed in to the 200ul Eppendorf tubes with 100μl of TLS. Tissue was mechanically homogenized, vortexed and centrifuged at 4°C and 5000 g for 30min. After isolation, the supernatant was collected, and the protein concentration measured using BCA assay. The tissue lysates were placed at −80°C until use.

#### Electrophoresis, western blot and immunoprecipitation

To confirm the synaptic changes after enzymatic or genetic manipulation of PNNs and hibernation like state. Series of pre and postsynaptic markers were measured. To separate proteins the Tris-glycine 4-15% precast gel on Biorad miniprotean aperture was used. Transfer was done on PVD membrane. Applied antibodies against postsynaptic markers: PSD95 (1:1000, Abcam, Cambridge, UK), GAD65/67 (1:3000, Abcam). Applied antibodies against presynaptic markers: vGLUT1 (1:1000, SYSY), vGAT (1:1000, Synaptic Systems), SNAP25 (1:1000, Synaptic Systems). Secondary antibodies Goat anti rabbit-HRP (1:15000, Abcam), Rabbit anti mouse –HRP (1:15000, Abcam) and goat anti-guinea pig-HRP (1:5000, Abcam) were used to visualize proteins of interest.

### Behavioral tests

#### Morris water maze

Morris water maze test for long term memory was applied to determine the impact of ChABC or local Acan removal in the hippocampus in model of AMP mediated hibernation like state synapse retraction. Interval of 60 s was given for animal to reach the target platform (at least 0.5 s interval at the target zone). When not reaching it, the value was automatically estimated as 60 s. Training period was for 5 consecutive days, four trials per day, each from different side of the pool (West, North, East and South). The order of release sites was changed every day to prevent scheme orientation. Two visible permanent cues were present. First day of training period mice were placed in to the MWM swimming pool with visible target platform. From the second day of training period the platform was hidden under the level of water (approx. 0.5 cm). Two weeks after training period, including stereotaxic injections of enzymatic treatment and hibernation like state protocol, the mice started experimental MWM period. Again, five consecutive days of four trials per day from four distinct locations.

#### Probe test

Probe test trials were applied to define the current state of memory of MWM task, and by comparison with the previous time points the ability to re-/learn the target position. The target platform was removed, and animals were placed from east entrance in to the pool for 60 s trajectory analysis. Four probe test trials held on day 1 (naïve animals), 6 (trained animals), 22 (memory retention test), 27 (re-learning ability.) were used to determine animal progress.

#### Y-Maze

Spontaneous alternation test in Y-maze was used at the end of experimental period to confirm no pathological changes in explorative behavior in new, experiment non-related arena. Mice were placed in the middle of triangular platform and the trajectory and entrance in the three distinct target zones were measured. When animal first reached one of the arms the error counting begun. Scoring; 1 point - reaching new arm, 2 points - reaching the arm, which was visited prior the latest, 3 points - reaching the latest arm again. The average error and the general activity in the maze (frequency of all arms visiting) was measured.

### Statistic

Using Sigma-stat software the study data were evaluated. For statistical evaluation of behavioral data, where the same subjects have been continuously assessed, the two-way repeated measurement ANOVA test was used. For confocal microscopy measurements, FIB-SEM microscopy and proteomic analysis (Western blot, WES), where treatment and time factor were considered, the two-way ANOVA was applied. Student-Neuman-Keuls post hoc multiple comparison test was used to study detail interactions, Holms-Sidak test was used to compere experimental data to control. Data presented in the graphs are expressed as arithmetic mean, with standard error of mean intervals. The normality of the tested values and statistical significance of the differences among groups in the text are described by F (H, q) values and p values, respectively. In the text and graphs when p≈ 0.05-# p<0.05 - *, p<0.01-**, p<0.001-***.

---

**Figure S1.**
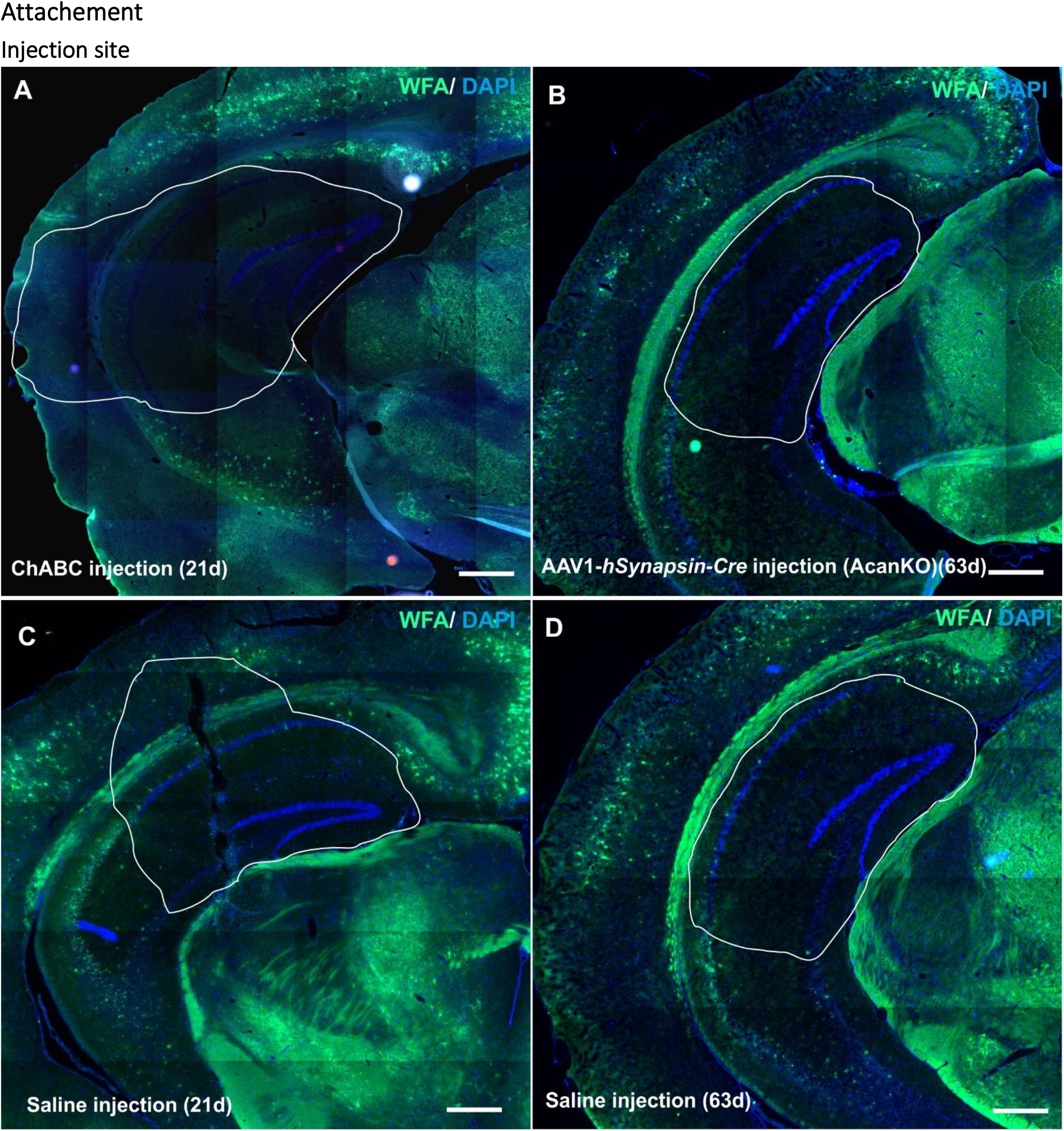
Injection site of ChABC (A,21 days after injection) and AAV1-*hSynapsin-Cre* AcanKO (B, 63 days after injection) at the end of the study (Day 28, Scheme1). Control injection sites showing no PNN digestion (WT-Saline (C), floxP - saline (D)). Stained on WFA (green) and DAPI (blue). Scale bar 500um.

**Figure S2.**
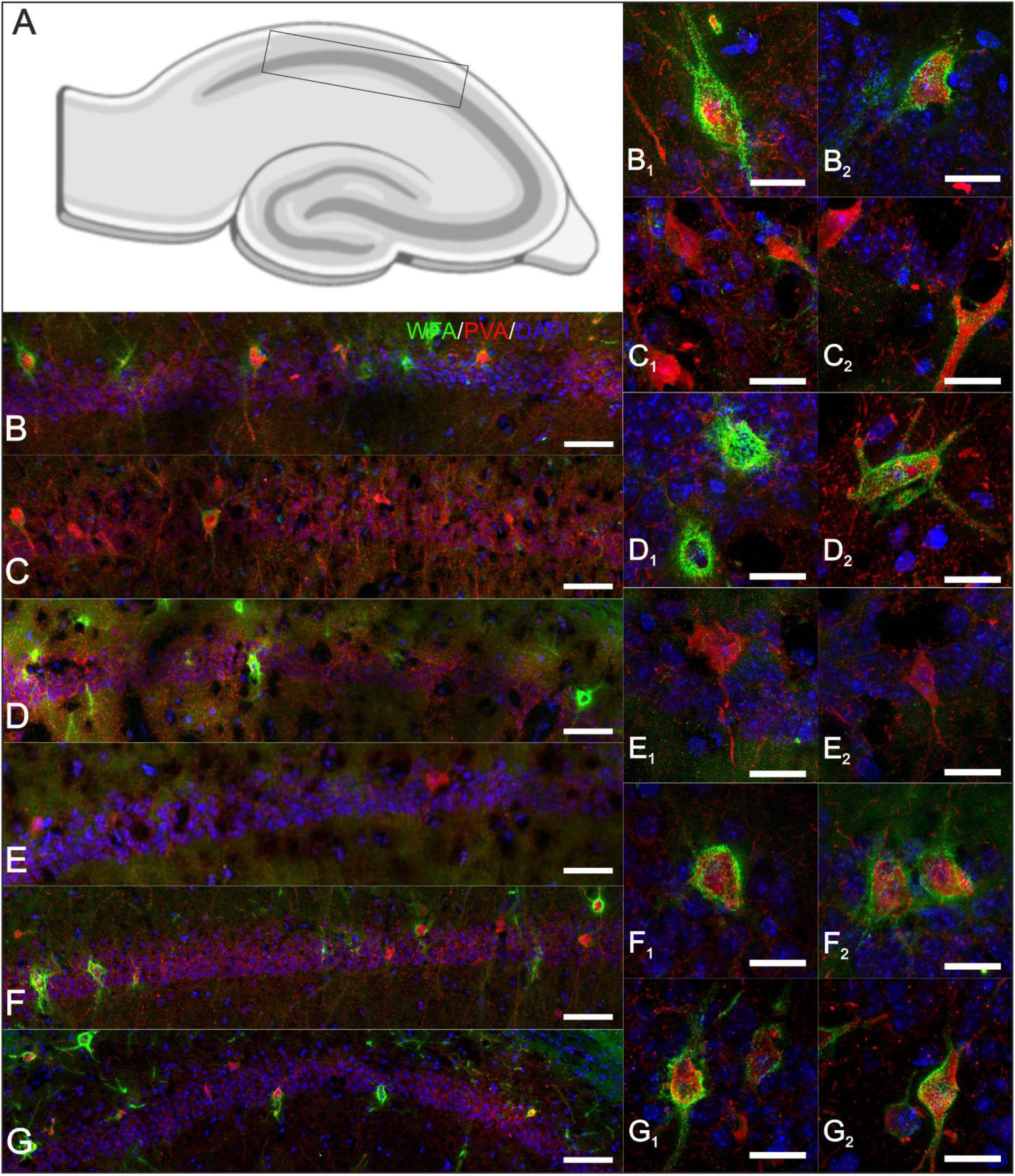
Detail image of PNN disintegration in the CA1 are of hippocampus (Illustrated location, A) of enzymatically treated (ChABC C, detail C1,2) or AcanKO animals (E, detail E1,2) at day 8, before HLS. In CA1 area of saline treated animals no sign of disintegration was observed (Saline group B, detail B1,2, floxP group, D detail D1,2). Intact WFA positive PNNS were observed in saline treated WT (F, Detail F1, 2) or floxP (G, detail G1, 2) mice that underwent HLS procedure. Samples were taken immediately after synapse withdrawal incubation period when BT was 16°C. WFA (Green) Parvalbumin (Red) DAPI (Blue). Scale bar 100μm (A - F), 10μm (A 1, 2-F 1, 2).

**Figure S3.**
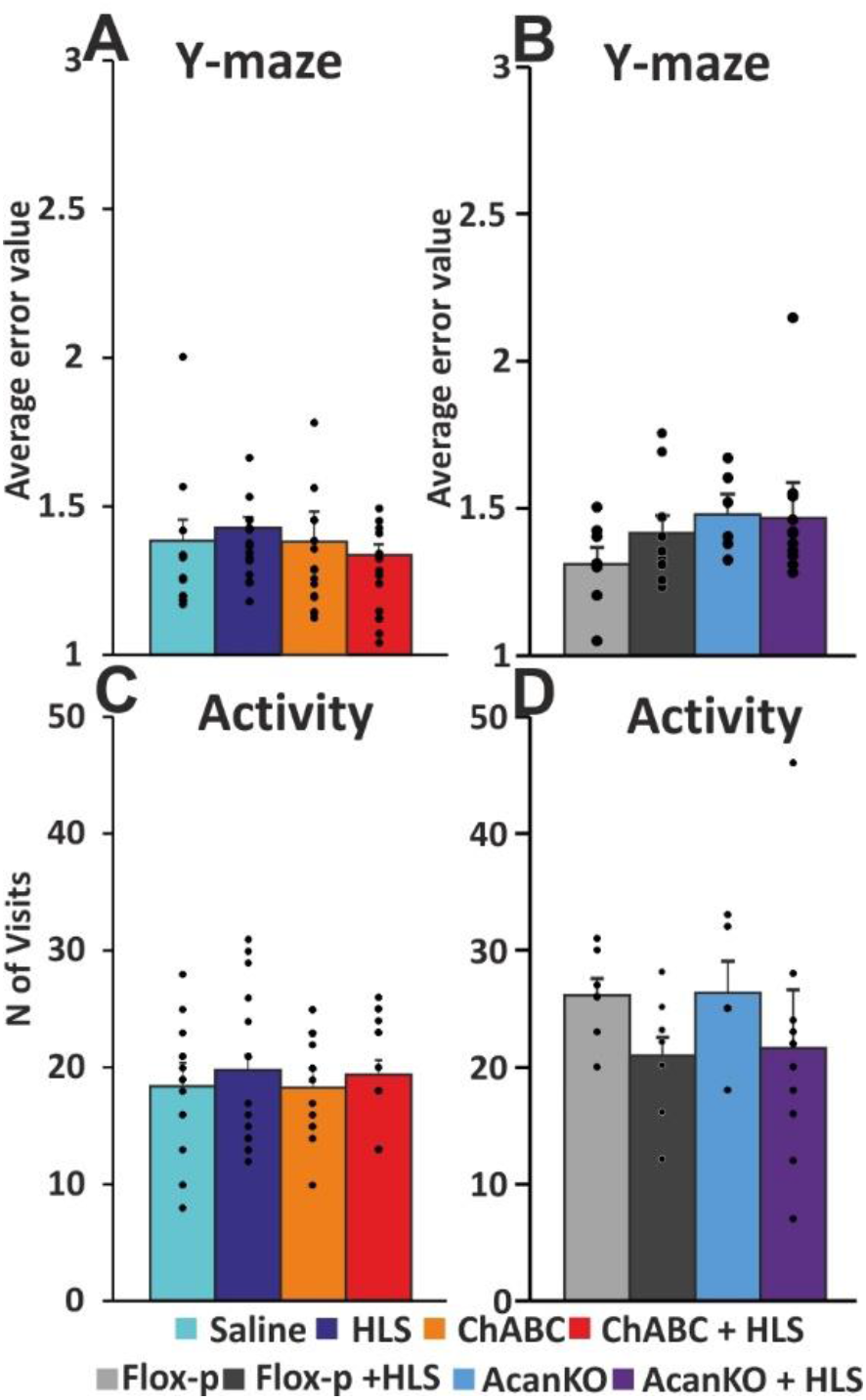
Spontaneous alternation test did not show any significant changes in working memory or general activity in the maze between treatment groups. Only a mild decrease in activity and increase error behaviour was observed in HLS groups

**Statistic table to figure 1.**
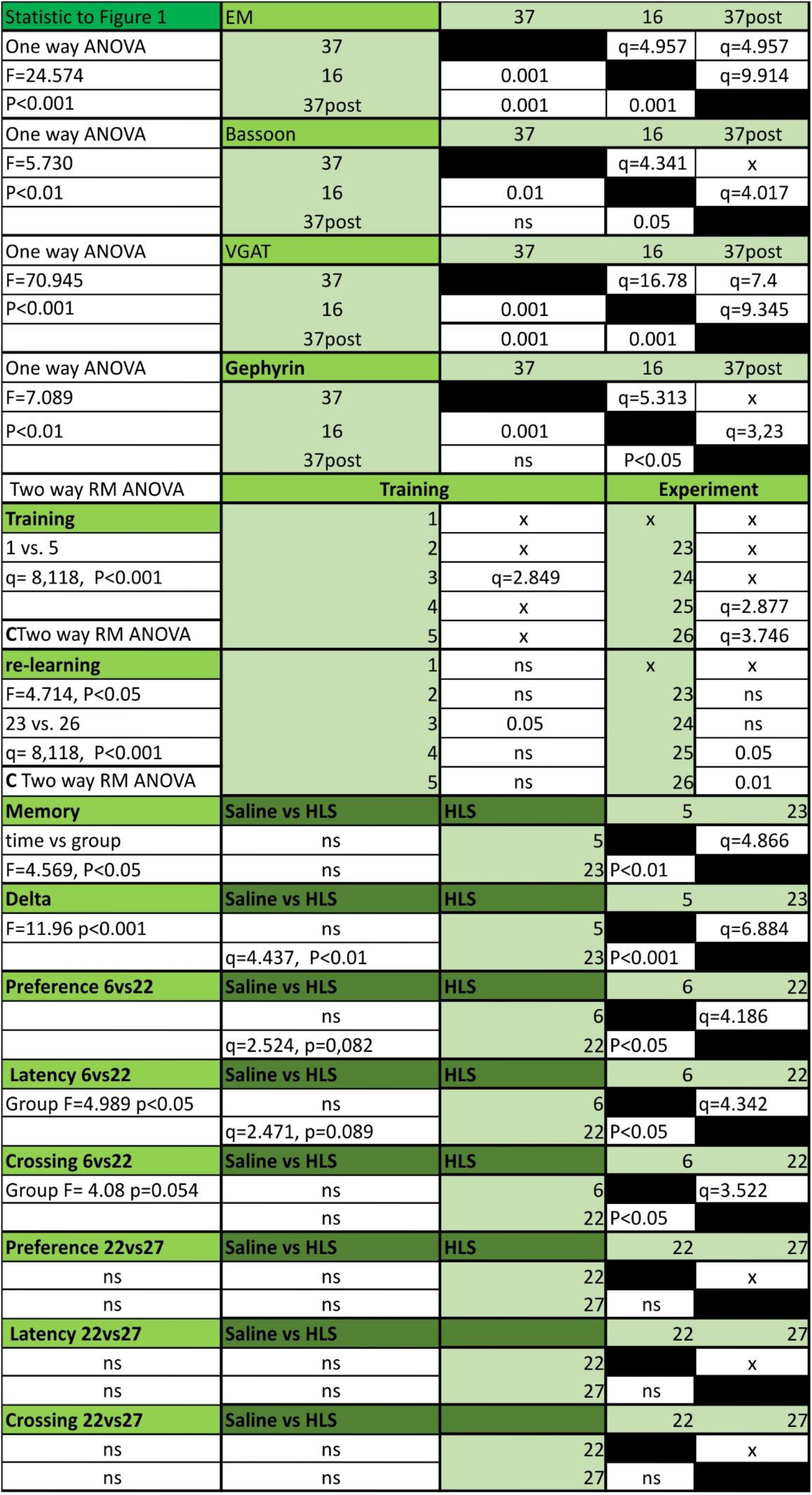

**Statistic table to figure 2.**
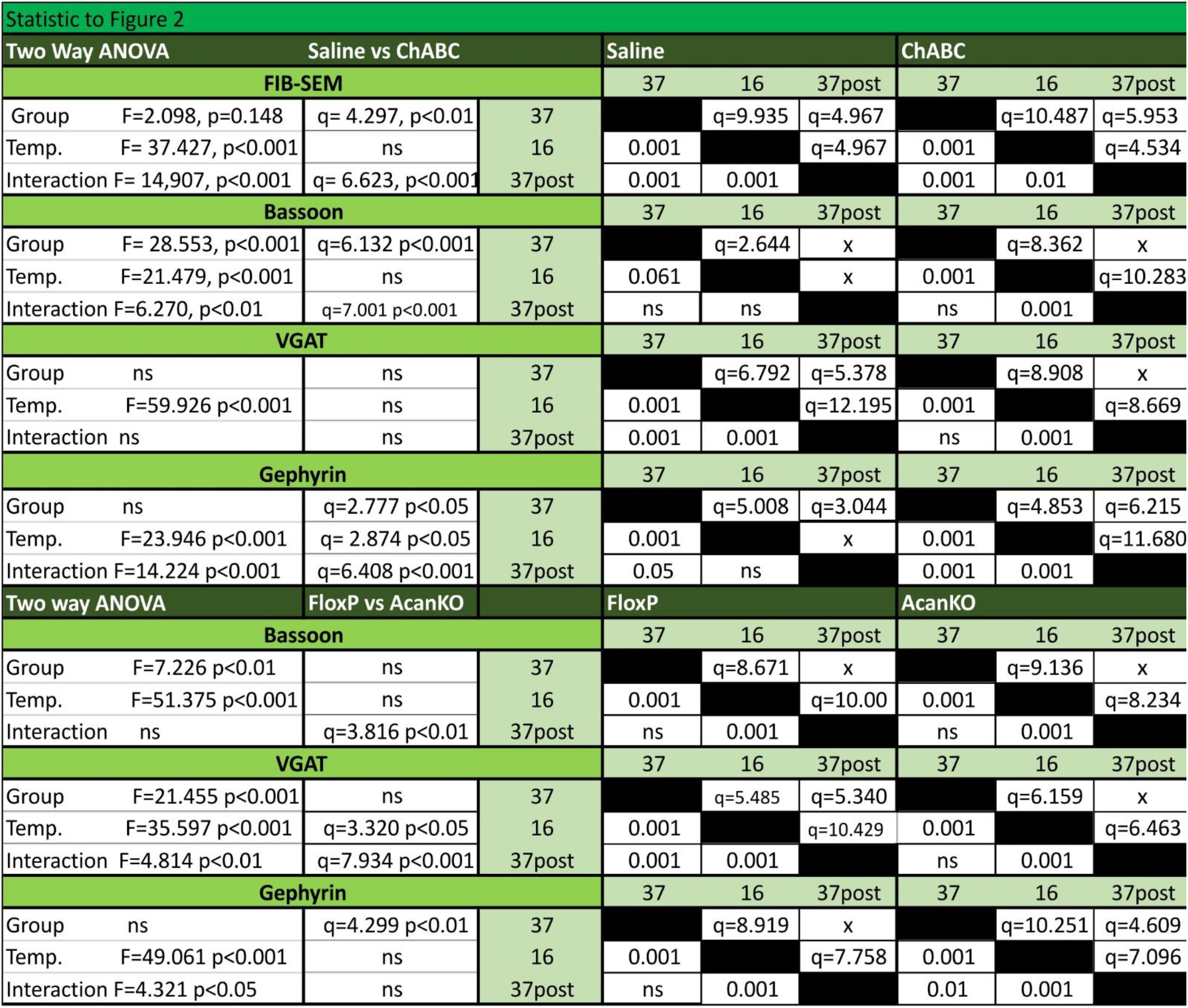

**Statistic table to figure 3.**

Statistic table to figure 2.
| Statistic to Figure 3 | One Way ANOVA |  |  |  |
| --- | --- | --- | --- | --- |
| **Bassoon** | F=12.065 p<0.001 |  |  |  |
|  | Saline | HLS | ChABC | ChABC +HLS |
| Saline |  | x | q=6.048 | x |
| HLS | ns |  | q=8.255 | q=3.605 |
| ChABC | 0.001 | 0.001 |  | q= 4.648 |
| ChABC + HLS | ns | 0.05 | 0.01 |  |
| **VGAT** | F=5.909 p<0.001 |  |  |  |
|  | Saline | HLS | ChABC | ChABC +HLS |
| Saline |  | q= 4.144 | q= 4.945 | q= 5.147 |
| HLS | 0.01 |  | x | x |
| ChABC | 0.01 | ns |  | x |
| ChABC + HLS | 0.01 | ns | ns |  |
| **Gephyrin** | F=6.804 p<0.001 |  |  |  |
|  | Saline | HLS | ChABC | ChABC +HLS |
| Saline |  | x | q=3.900 | x |
| HLS | ns |  | q=6.320 | x |
| ChABC | 0.01 | 0.001 |  | q=4.203 |
| ChABC + HLS | ns | ns | 0.05 |  |
| **Bassoon** | F= 5.905 p=0.001 |  |  |  |
|  | FloxP | FloxP + HLS | AcanKO | AcanKO +HLS |
| FloxP |  | q=4.077 | x | x |
| FloxP + HLS | 0.01 |  | q= 5.636 | q= 4.507 |
| AcanKO | ns | 0.001 |  | x |
| AcanKO +HLS | ns | 0.01 | ns |  |
| **VGAT** | F= 18.753 p<0.001 |  |  |  |
|  | FloxP | FloxP + HLS | AcanKO | AcanKO +HLS |
| FloxP |  | q=7.256 | q= 2.457 | q=9.734 |
| FloxP + HLS | 0.001 |  | q=4.206 | q=2.664 |
| AcanKO | 0.085 | 0.01 |  | q= 6.511 |
| AcanKO +HLS | 0.001 | 0.062 | 0.001 |  |
| **Gephyrin** | ns |  |  |  |
|  | FloxP | FloxP + HLS | AcanKO | AcanKO +HLS |
| FloxP |  | x | x | x |
| FloxP + HLS | ns |  | x | x |
| AcanKO | ns | ns |  | x |
| AcanKO +HLS | ns | ns | ns |  |

**Statistic table to figure 4.**
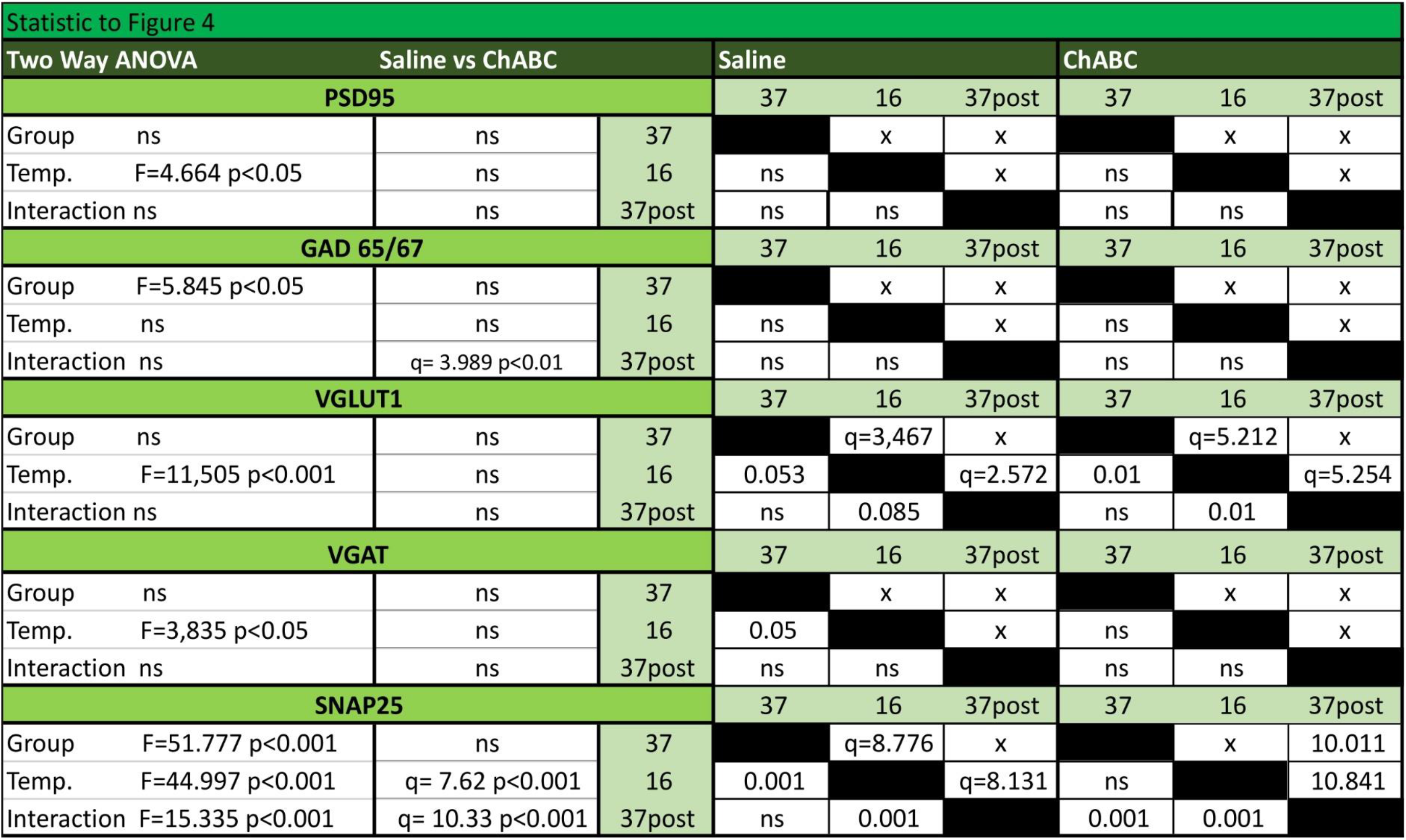

**Statistic table to figure 5.**

Statistic table to figure 4.
| Statistic to Figure 5 | One Way ANOVA |  |  |  |
| --- | --- | --- | --- | --- |
| **PSD95** | ns |  |  |  |
|  | Saline | HLS | ChABC | ChABC +HLS |
| Saline |  | x | x | x |
| HLS | ns |  | x | x |
| ChABC | ns | ns |  | x |
| ChABC + HLS | ns | ns | ns |  |
| **GAD65/67** | F=2.754 p=0.06 |  |  |  |
|  | Saline | HLS | ChABC | ChABC +HLS |
| Saline |  | q=3.690 | x | x |
| HLS | 0.064 |  | q=3.127 | x |
| ChABC | ns | 0.086 |  | x |
| ChABC + HLS | ns | ns | ns |  |
| **VGLUT1** | F=6.701 p<0.001 |  |  |  |
|  | Saline | HLS | ChABC | ChABC +HLS |
| Saline |  | q=3.374 | q=6.194 | x |
| HLS | 0.059 |  | q=3.155 | x |
| ChABC | 0.001 | 0.05 |  | q=4.15 |
| ChABC + HLS | ns | ns | 0.05 |  |
| **VGAT** | ns |  |  |  |
|  | Saline | HLS | ChABC | ChABC +HLS |
| Saline |  | x | q=2.635 | x |
| HLS | ns |  | x | x |
| ChABC | 0.072 | ns |  | x |
| ChABC + HLS | ns | ns | ns |  |
| **SNAP25** | F=4.503 p<0.01 |  |  |  |
|  | Saline | HLS | ChABC | ChABC +HLS |
| Saline |  | x | q=4.367 | q=3.531 |
| HLS | ns |  | q=3.694 | q=2.812 |
| ChABC | 0.05 | 0.05 |  | x |
| ChABC + HLS | 0.05 | 0.056 | ns |  |

**Statistic table to figure 6.**
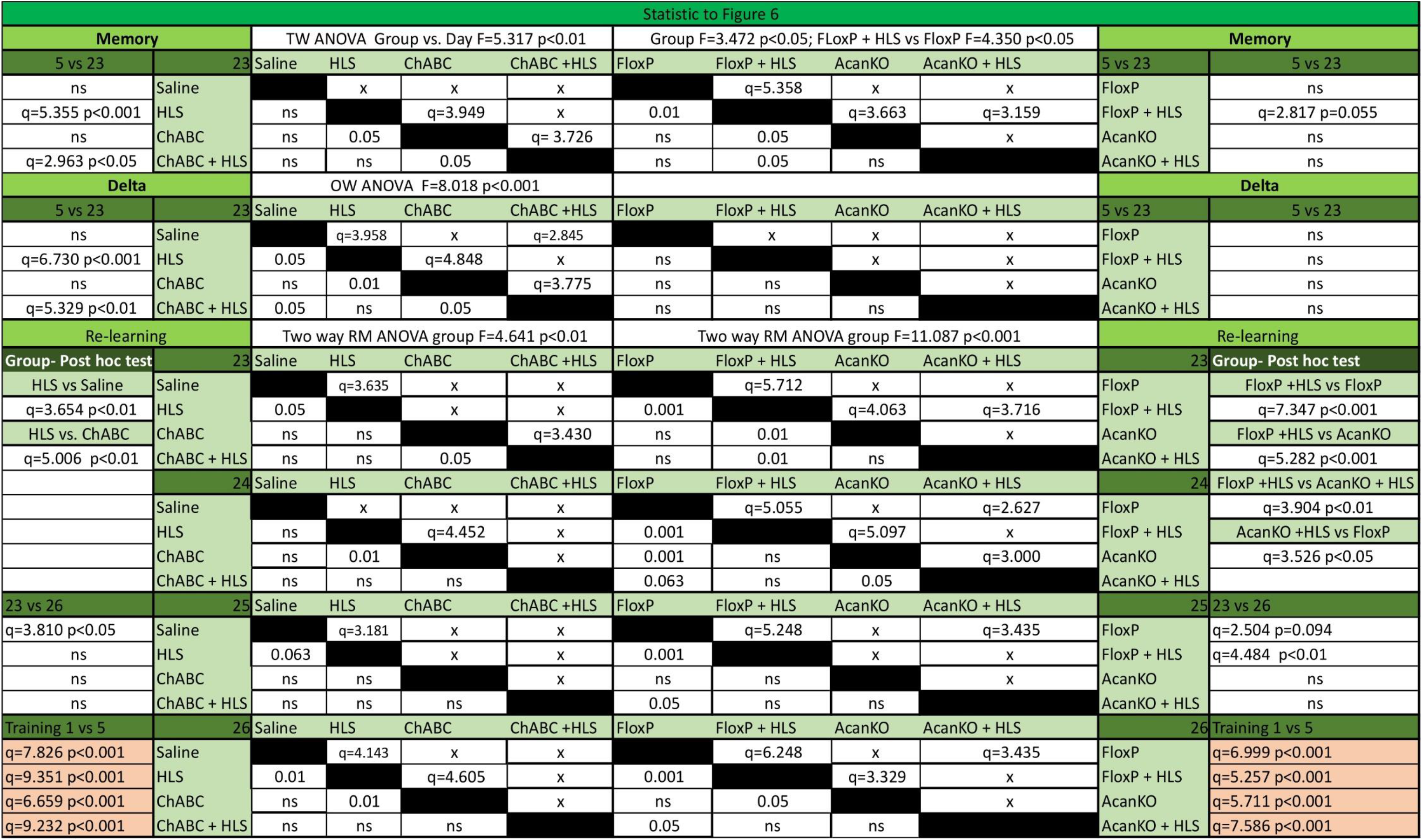

**Statistic table to figure 7.**

Statistic table to figure 6.
| Statistic to figure 7 |  |  |  |
| --- | --- | --- | --- |
| Two way ANOVA |  |  |  |
| 6 vs 22 |  |  |  |
|  | Preference |  | Group vs Day F=3.015 p<0.05 |
| ns | Saline | FloxP | ns |
| q=4,412 p<0.001 | HLS | FloxP + HLS | ns |
| ns | ChABC | AcanKO | q=2.810 p=0.056 |
| q=2.671 p=0.076 | ChABC + HLS | AcanKO + HLS | q=3.017 p<0.05 |
| Crossing |  |  |  |
| ns | Saline | FloxP | ns |
| ns | HLS | FloxP + HLS | q=2.632 p=0.073 |
| q=4.346 p<0.01 | ChABC | AcanKO | ns |
| ns | ChABC + HLS | AcanKO + HLS | ns |
| Latency |  |  |  |
| ns | Saline | FloxP | ns |
| q=4.240 p<0.01 | HLS | FloxP + HLS | q=2.752 p=0.061 |
| ns | ChABC | AcanKO | ns |
| ns | ChABC + HLS | AcanKO + HLS | ns |
| Quadrant |  |  |  |
| ns | Saline | FloxP | ns |
| q=2.749 p=0.069 | HLS | FloxP + HLS | ns |
| q=2.99 p=0.05 | ChABC | AcanKO | ns |
| q=6.004 p<0.001 | ChABC + HLS | AcanKO + HLS | ns |

**Statistic table to figure 8.**
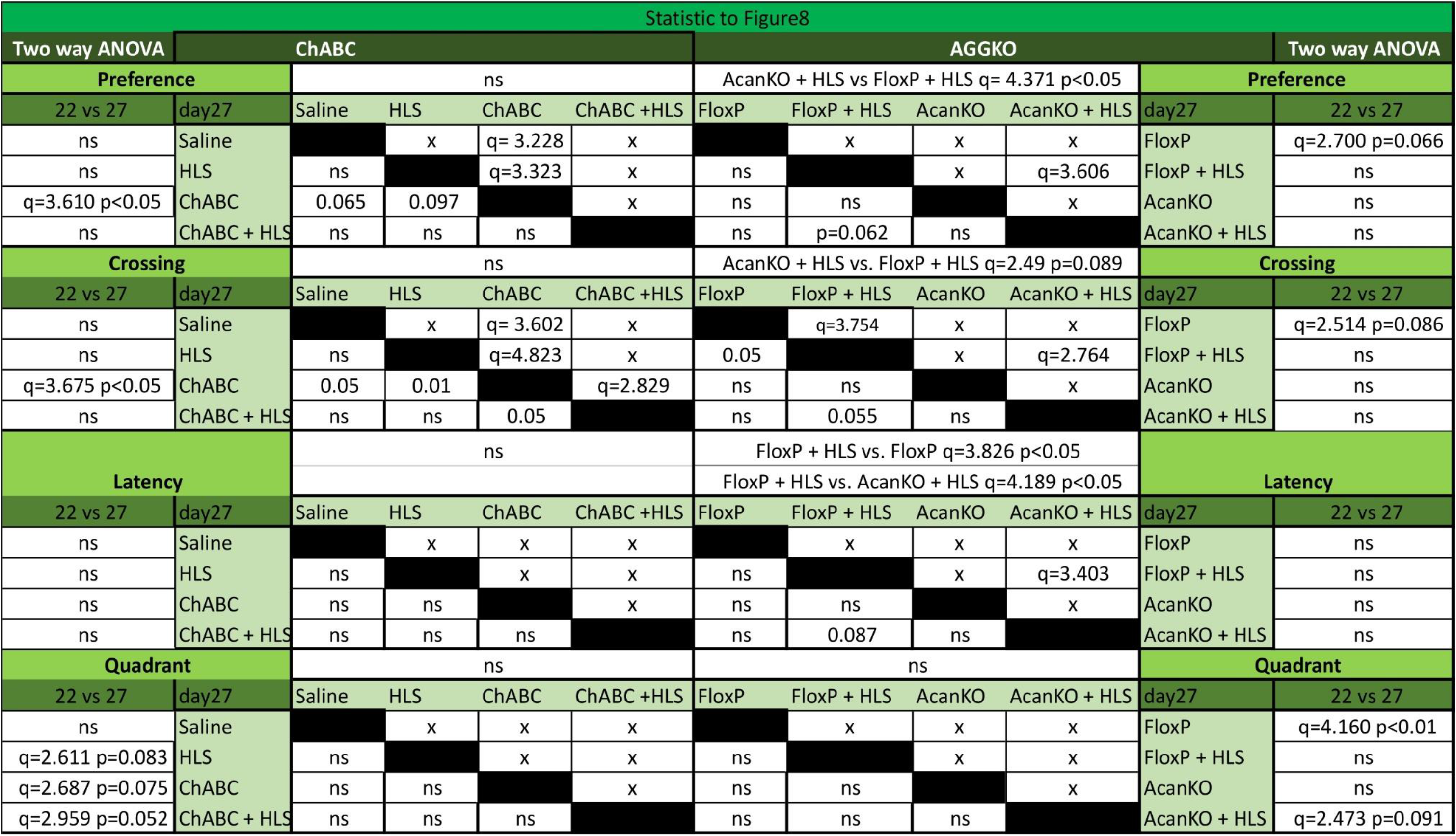

